# Time-varying co-activity and connectivity of the somato-cognitive-action-network in densely sampled brains

**DOI:** 10.1101/2025.07.16.660604

**Authors:** Richard F. Betzel

## Abstract

The somato-cognitive-action-network (SCAN) is a recently discovered brain network. SCAN interdigitates somatomotor networks and is functionally connected to the cingulo-opercular/action-mode network. SCAN is therefore thought to play a role in motor/action planning. This view of SCAN, however, is based on its “static” functional connectivity–i.e. FC estimated using data pooled over an entire scan session or dataset. However, this approach necessarily overlooks changes in network activity and connectivity that occurs over shorter timescales. In this report, we extend analyses of SCAN’s static architecture, demonstrating that, at the whole-brain level, SCAN vertices exhibit overlapping network membership, such that, while they form a cohesive sub-network, they may also couple transiently and dynamically with other networks. We examine these potential co-activations by focusing on SCAN dynamics and through the identification of distinct coupling modes – co-activation patterns (CAPs). We show that CAPs differentially contribute to SCAN’s static architecture and that their frequency varies across following limb immobilization.

## INTRODUCTION

Multi-modal imaging studies have found that the human brain can be meaningfully divided into sub-systems (henceforth referred to as functional networks) [1–4]. Functional networks tend to be dissociable from one another on the basis of their function [5], architectonics [6], connectivity [1], and/or topographical representation [7]. The spatial boundaries between networks are evident, even in the absence of explicit task instructions– i.e. during the so-called “resting state”– as communities and clusters in intrinsic functional connectivity (FC) [8].

Despite the intense interest in network mapping, there still exists controversies–e.g. the appropriate nomenclature for referring to networks [9, 10]. Additionally, new experimental and analytic approaches occasionally provide evidence of altogether new networks. Recently, using resting state fMRI data from densely-sampled individuals, Gordon et al. [11] discovered the somato-cognitive-action network (SCAN). SCAN is composed of six approximately bilaterally symmetric “inter-effector” regions that interdigitate somatomotor networks along primary motor cortex, challenging conventional perspectives on the organization of somatotopic representations in M1 [12]. SCAN also included secondary cortical components in supplementary motor area and anterior cingulate, as well as sub-cortical and cerebellar elements, with strong functional connectivity to putamen and thalamus.

SCAN’s connectivity positions it as a mediator between regions in the cingulo-opercular (or “action-mode” [13]) network that supports goal-directed cognition and somatomotor effector regions [14, 15]. This has led to speculation that SCAN may represent an ideal target for neuromodulation, especially in the service of movement disorders–e.g. Parkinson’s disease [16, 17]. More generally, however, and owing to its relative recent discovery, SCAN’s architectural features, including its temporal stability, have not been fully explored.

The aim of this paper is simply to fill in some of these gaps. It is structured not as a test of any specific set of hypotheses; rather the goal of this paper is more modest, seeking to catalog potentially interesting and previously undisclosed features of SCAN. While some of features are time-invariant–e.g. derived from time-averaged and so-called “static” connectivity– the broader focus is on the spatiotemporal structure of SCAN–i.e. how the activity of inter-effector regions waxes and wanes over short timescales, how the rest of cortex behaves during those fluctuations, how these patterns map onto moment-to-moment changes in SCAN connectivity, and ultimately how these dynamics contribute to SCAN’s static connectivity.

This general topic–time-varying activity and connectivity [18]–is central to network and cognitive neuroscience and, though not without controversy [18–20], has been used in the past to help elucidate the principles by which brain networks reconfigure in response to complex stimuli [21], spontaneously in the absence of explicit task instruction [22–27], and how those changes may be linked to behavioral, physiological, and demographic features [28–31].

This manuscript make several contributions over the course of the following sections. First, we show that SCAN FC is not homogeneous; rather, each inter-effector region maintains a distinct connectivity profiles with dissociable network targets. Second, we show that SCAN vertices maintain a plurality of network assignments, dually positioning SCAN as a separate network, but also as an integrative hub of network overlap–a module sitting atop other modules [32, 33]. Third, we show that frames in which SCAN exhibits high-amplitude activity can be partitioned into six co-activation patterns (CAPs) that recur across time, both within and across imaging sessions, and whose topography is broadly shared across individuals. Fourth, we show that after transforming CAPs into seed connectivity maps, they individually diverge from but collectively explain SCAN’s static FC profile, positing dynamic origins of SCAN connectivity. Fifth, we show that many, but not all, of the CAPs estimated at the group level are preserved within individuals. Sixth (and finally) we show that normalized CAP rate is modulated following limb immobilization.

In summary, this paper makes no singular contribution, but it identifies several previously undisclosed features of SCAN that might serve as useful targets in future applied studies.

## RESULTS

Here, we analyzed imaging data from two densely sampled cohorts. The first came from the so-called “Midnight Scan Club” [34] and consisted of 10 participants each scanned over the course of 10 imaging sessions. Each session included 30 minutes of resting state data, yielding (prior to motion censoring) 5 hours of imaging data per participant. We also analyzed data from the casting-induced plasticity dataset (henceforth referred to as the “Casting” dataset), which consisted of three individuals who underwent between 42 and 64 daily 30-minute resting state scans prior to, during, and following a limb-immobilization procedure in which they wore a cast over their right upper arm [35–37]. This dataset yielded between 23.5 and 31.5 hours of data per participant. For acquisition and processing details, see **Materials and Methods**. In principle, we could have also examined larger, shallower samples, e.g. imaging data from Human Connectome Project [38]. However, previous work has noted that success in detecting SCAN within individuals is partly determined by volume of data. The same is true (and perhaps more so) for the accurate characterization of brains’ time-varying profiles [20]. For these reasons, we opted to focus on dense-sampling datasets.

### Mapping the somato-cognitive action network with probabilistic and temporal template-matching

Prior to Gordon et al. [11], SCAN had gone undetected, despite more than a decade of intense interest in brain network mapping [1, 2, 8, 40]. This is due, at least in part, to historical emphasis on “group average” brains and generation of correspondingly representative network maps. Although a common procedure in non-invasive brain network mapping [1, 2, 41], averaging connectivity data across many brains effectively smooths the boundaries of network maps due to individualized variation in their topography, making it difficult to resolve fine-scale features. Consequently, SCAN–comprised of small, spatially disjoint inter-effector regions–is largely absent from contemporary network atlases [42], necessitating, for the time being, its manual detection and labeling in new imaging datasets.

Template matching [43] is a framework that accelerates network detection by comparing a participant’s imaging data (usually dense connectivity) against a set of network “templates” – i.e. seed connectivity maps spanning an atomized group of pre-defined networks. The connectivity of each vertex or voxel is compared against that of each template and assigned the label of the template to which it is maximally similar. This procedure is sometimes followed by a secondary cleaning step in which small and isolated network components are reassigned to larger neighboring networks.

Here, we used a semi-automated and probabilistic variant of template matching to assign network labels to each surface vertex based on their temporal features. Briefly, we used a probabilistic network atlas to create a weighted sum of vertex time courses for each network; we termed these network-representative time courses as *time series templates* [39]. Then, we computed the correlation of vertex time courses with each template, and assigned vertices the network label to which they were maximally correlated (see Fig. 2). Note that, unlike the standard template matching procedure, which is based on seed connectivity, ours is based on topography. Consequently, our approach does not require matching to a pre-calculated connectivity template, whose elements may be influenced by processing decisions. Rather, our procedure uses a population-level spatial prior to create time series templates tuned to each participant.

In explorations, we find that network labels stabilize with relatively few frames worth of data (see Fig. S1 for convergence rates from participants in the “casting” study). We note that, in principle, the spatial priors could also be used to estimate personalized seed connectivity templates–e.g. by created weighted sums of vertices from `.dconn`s; however, we opted to work with the `.dtseries` data, as it obviates the computationally expensive operation of generating vertex × vertex connectivity matrices.

In Fig. 1a, we show the MSC modal assignments for each vertex as well as the SCAN seed connectivity map (averaged over all SCAN vertices) to all other surface vertices Fig. 1b. We annotated this map to highlight three pairs of approximately bilaterally symmetric inter-effector nodes that comprised the canonical cortical component of SCAN. Though note that, as in Gordon et al. [11], we found circumscribed regions within anterior cingulate cortex that, here, were assigned to the cingulo-opercular/action-mode network [13], but could also be included as members of an extended SCAN based on their resting-state FC.

**Figure 1.**
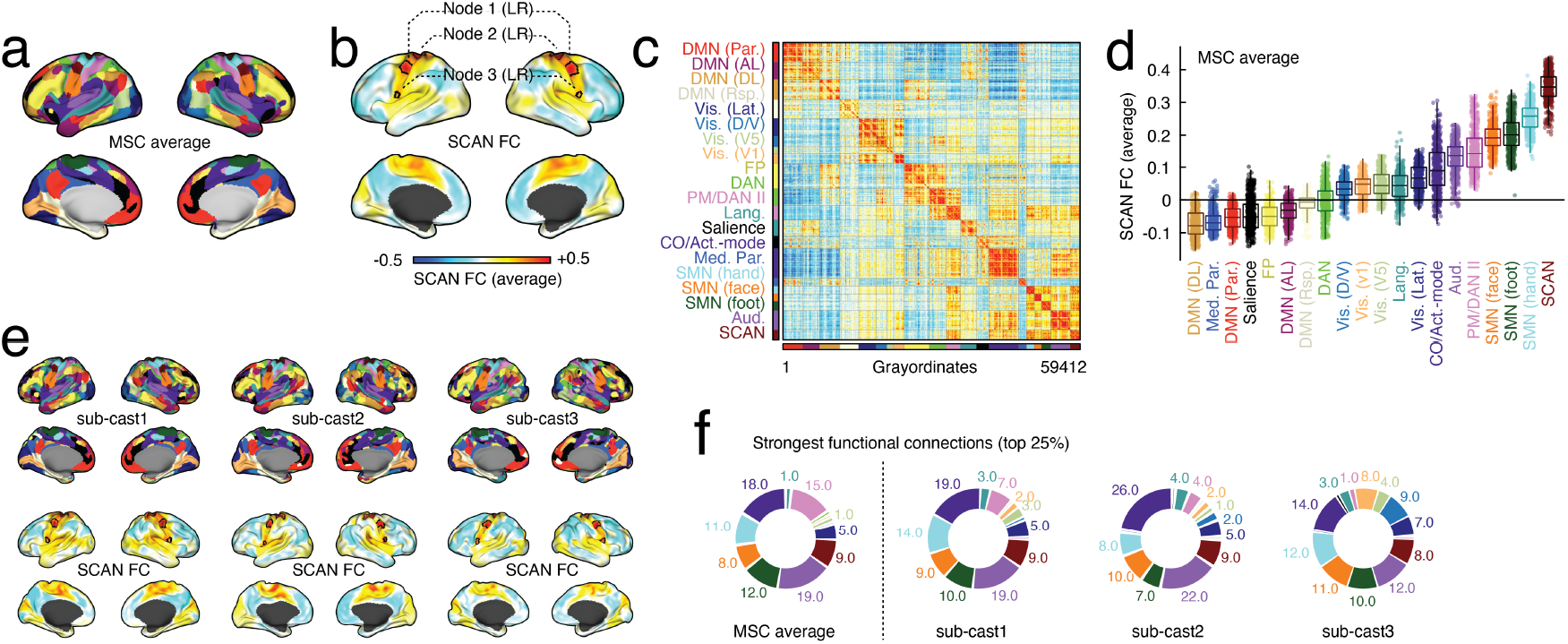
Somato-cognitive-action-network correlation structure. (*a*) Brain networks (systems) estimated using probabilistic and temporal template matching after pooling dense time series data from 10 participants in the MSC dataset. (*b*) SCAN is composed of three core nodes, highlighted here (black outlines) along with the FC seed map averaged over all SCAN vertices. (*c*) Whole-brain dense connectivity matrix with scan listed as the final network. (*d*) Mean FC weights (correlations) from SCAN to all other networks. (*e*) Single-participant network maps and SCAN seed FC for casting dataset. (*f*) System composition of strongest connections (top 25%) for MSC-averaged and single-participant data.

We recapitulated prior work, showing that SCAN maintained strong connectivity to other networks (Fig. 1c), notably its spatially proximal neighbors, which included all labeled sub-components of the somatomotor network (hand, foot, face), along with the pre-motor, auditory, and cingulo-opercular/action-mode networks (Fig. 1d). Although, like other networks, we observed variation across participants in terms of SCAN topography (see Fig. 1e for network maps from casting participants), the overall composition of SCAN’s strongest FC, at least in terms of networks, was largely consistent across individuals (Fig. 1f).

### Variation in FC across SCAN vertices

In the previous section, we described six regions (three bilaterally symmetric pairs) that comprised the cortical component of the canonical SCAN. Like other networks, ascribing a singular label to SCAN vertices can falsely give the impression that SCAN is a homogeneous object and potentially obscure variation of connectivity profiles within SCAN. Here, rather than characterize the mean connectivity of SCAN to the rest of cerebral cortex, we examined heterogeneity of connectivity across the three node pairs.

To do this, we separately estimated whole-brain FC seed maps for each SCAN node. Due to strong hemispheric symmetry of seed maps, we combined the left and right analogs into node-specific maps (three pairs in total). We then quantified, for each map, its connectivity to six networks with the strongest FC to SCAN. These included the cingulo-opercular/action-mode, auditory, pre-motor/dorsal attention II, and all somato-motor networks.

We observed a consistent spatial gradient of FC in both MSC and casting datasets. Moving along the pre-central gyrus in a superior medial-to-inferolateral direction, we found increased FC of SCAN to the cingulo-opercular/action-mode, auditory (though weak), and somatomotor (face) networks (Fig. 3). Connectivity to somatomotor (foot) and pre-motor/dorsal attention II networks decreased, while connectivity to somatomotor (hand) exhibited an inverted u-shape.

**Figure 2.**
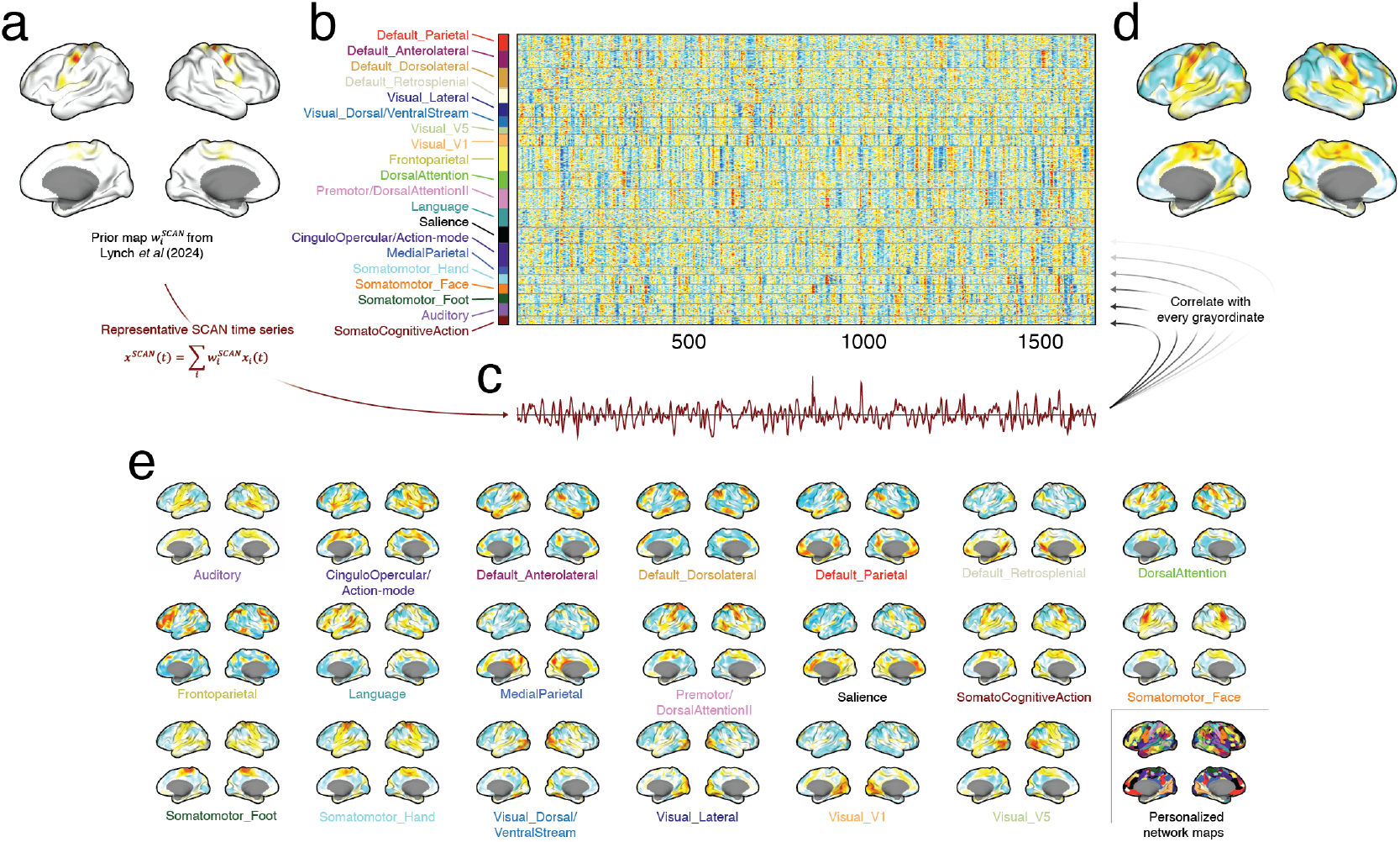
Probabilistic temporal template matching for defining personalized networks, including SCAN. This figure outlines the procedure used to obtain personalized network maps. (*a*) Using previously estimated network probability maps [39], we calculated a representative time series for each network – a template time series – as the sum of all vertex time series (see *b*), each weighted by the probability that it belongs to a given network at the population level. (*c*) As an example, we show the time series template for SCAN. (*d*) After time series templates are created, we correlate them with each vertex. We show the correlation of vertices with the SCAN template here. (*e*) Template correlation maps for 20 networks. We estimate the primary network assignment of each vertex based on which of the 20 templates it is maximally correlated. We show an example set of personalized network labels in the bottom right.

**Figure 3.**
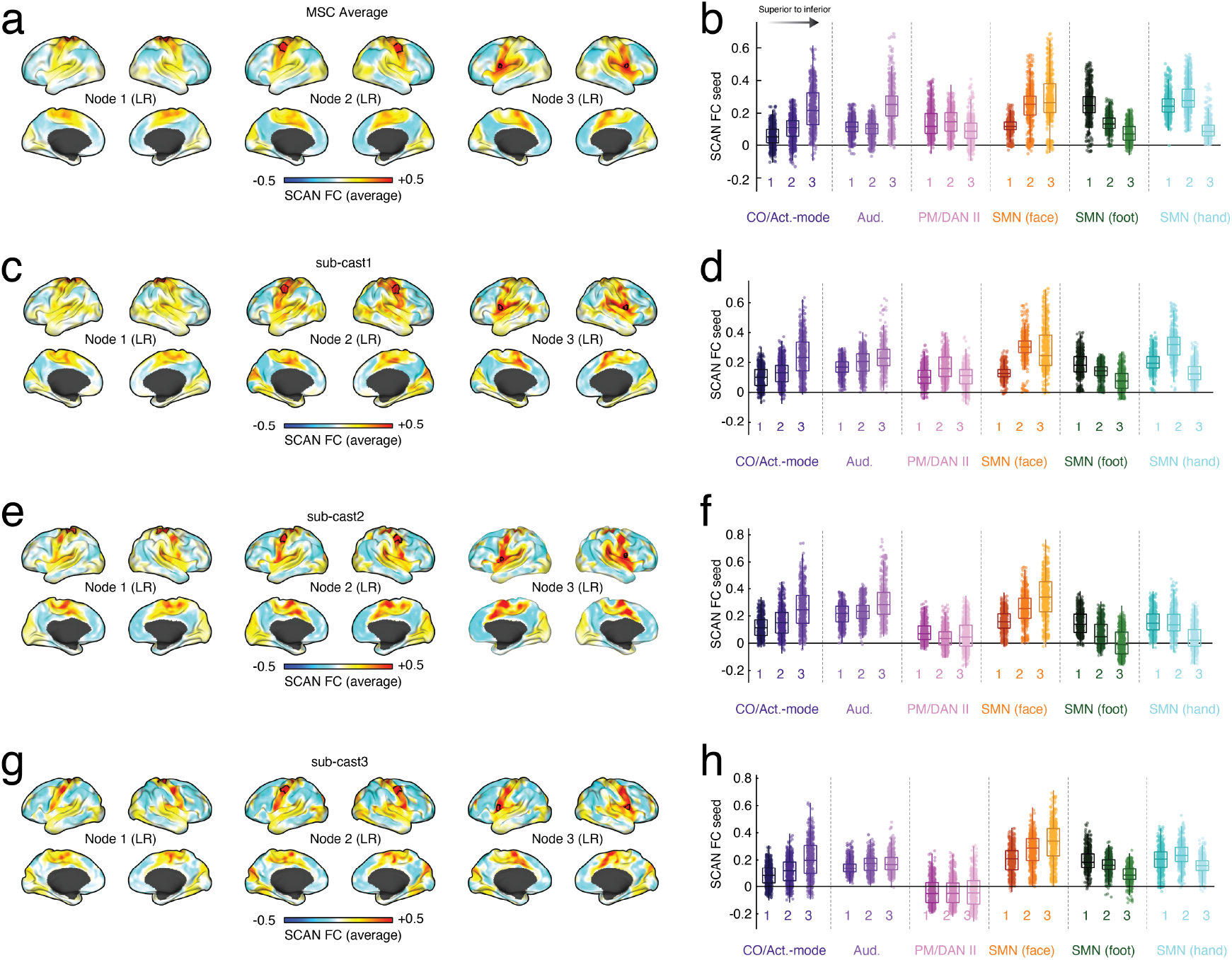
Within-SCAN FC spatial variability. SCAN is composed of six bilaterally symmetric nodes that can be grouped into three pairs. To this point, we have described their mean connectivity. Here, we examine the connectivity of each node pair independently. (*a*) FC seed maps for each node. (*b*) Boxplots for each nodes’ connections to other networks that comprise the somatomotor-action complex. Panels *a* and *b* show results for pooled MSC data. Panels *c*-*h* show analogous plots for the three participants in the casting study.

Collectively, these results suggest that SCAN, though treated as a uniform entity, nonetheless exhibits specific patterns of variation in FC across inter-effector regions.

### SCAN nodes are network hubs and integration sites for sensorimotor and action networks

Here, we used a variant of template matching to map SCAN at the single-subject level. Although this algorithm is a powerful approach for individualized network mapping, the “winner take all” step is a limitation and can return potentially misleading results. For instance, vertices with no clear correspondence to any network are nonetheless assigned a network label. Another, and potentially more interesting issue, concerns is the case where a vertex has strong correspondence to multiple networks (and could therefore potentially be classified as a “hub”). Despite their poly-affinity, these vertices would nonetheless be assigned a singular label corresponding to their top match; their “primary” network assignments. An alternative strategy, and one that possibly circumvents this issue, is to allow vertices to take on multiple network assignments, provided that their similarity to a given network template exceeds some minimum level, *r*_*min*_ (see Fig. 4a). This overlap score– the number of networks a vertex is assigned to–can then be interpreted as a measure of “hubness” and network-level integration.

**Figure 4.**
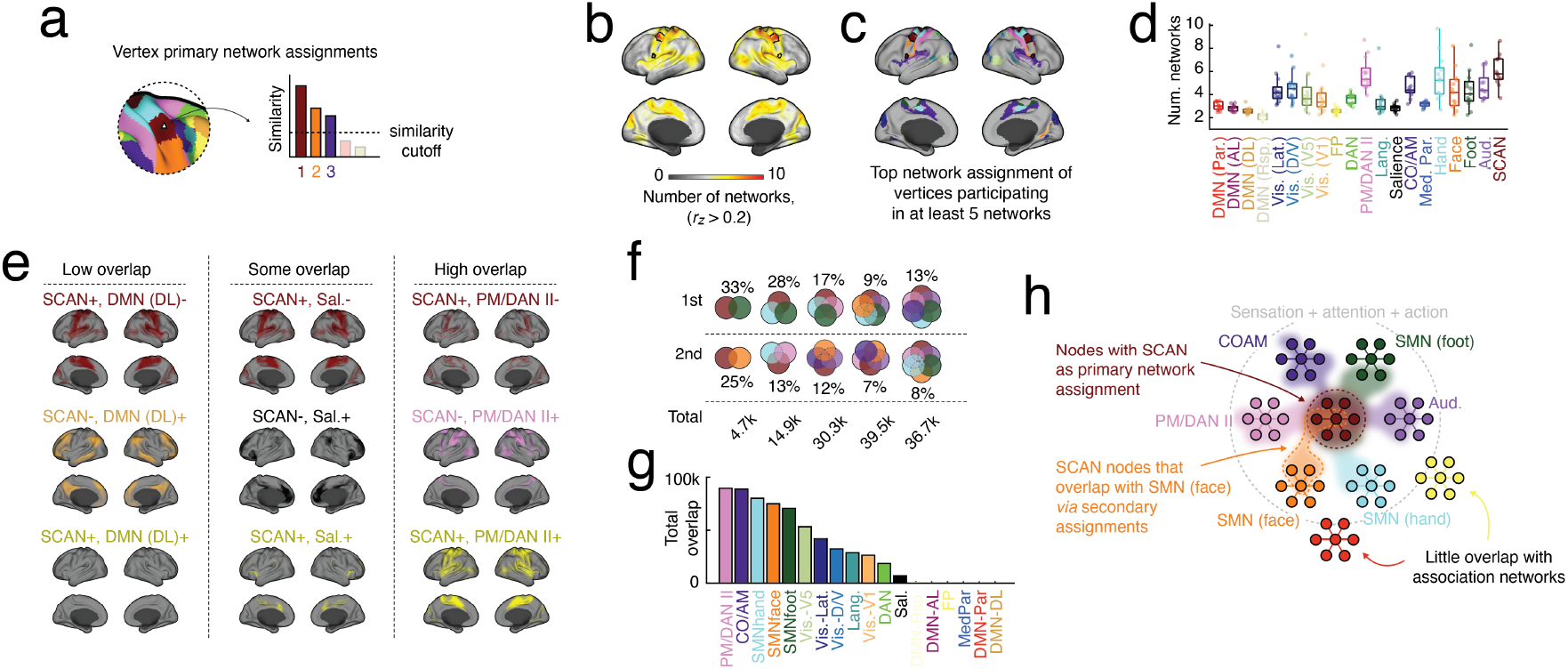
Secondary network assignments position SCAN vertices as an integrative, multi-network hub. Template matching algorithms rely on a “winner-take-all” step to assign vertices a network label. However, it can be the case, for a given vertex, that while there is a winner, the next best network match is a small *ε* below. In such a case, the secondary network assignment may be as meaningful as the first. We illustrate this example in *a*. Here, for each vertex, we count the number of networks with similarity > *r*_*min*_. We then estimate probabilistic maps for each network based not only on vertices’ primary network assignments, but any network satisfying the criterion, > *r*_*min*_. We can count the number of networks to which each vertex time course was correlated with above > *r*_*min*_ and average this number across participants. We show the result in *b* and in *c*, the modal network labels of vertices that participated in, on average, at least five networks. (*d*) Boxplot showing mean network count for all networks. Each point represents a data point from a single individual. We can also count how frequently specific *pairs* of networks overlap and represent those frequencies in anatomical space. For instance, SCAN and the dorsolateral component exhibit almost no overlap (*e*, left). The top plot shows vertices that were assigned to SCAN but not DMN (DL). The middle plot shows vertices assigned to DMN (DL) but not SCAN. The final plot shows the intersection–i.e. vertices assigned to both. Other networks exhibit modest amounts of overlap. See, for instance, SCAN and the salience network (*e*, middle). Others exhibited much greater amounts of overlap. See SCAN and PM/DAN II (*e*, right). (*f*) Focusing on SCAN, we identified vertices in which SCAN overlapped with exactly one, two, three, four, and five other networks. We show here the first and second most frequent pairing. For instance, if we were to restrict ourselves to vertices in which SCAN was supra-threshold with exactly one other network, we find that there are approximately 4,700 such vertices, in total and that roughly 1/3 of these pairings are SCAN with SMN (foot). Of the remaining pairs, 25% were SCAN with SMN (hand). Similarly, if we considered vertices where SCAN was paired with four other networks, we find that there were 39,500 such vertices, the top quintet included SCAN with all three SMN sub-networks along side the auditory network (9%) while the second most frequent quintet included SCAN and auditory, PM/DAN II, SMN (face), and cingulo-opercular/action-mode networks. (*g*) We also counted total overlap with SCAN, irrespective of network count per vertex. (*h*) Together, our findings suggest that SCAN overlaps considerably with networks associated with sensation, attention, and action. In contrast, SCAN exhibits little overlap with association networks, e.g. default mode or fronto-parietal.

Here, we calculated overlap scores for each participant based on their own dense connectivity data. We subsequently generated a composite map by averaging overlap scores across participants. Interestingly, we found the vertices with SCAN as their primary network assignment also tended to have the greatest overlap score (Fig. 4b,c,d; *t*-test, SCAN *versus* other networks; *p* < 0.05), a feature that may be related to the diversity of networks with which it shares a border (Fig. S3). Zooming out, the level of overlap was elevated in the somatomotor-action complex: a collection of networks that included all three somatomotor networks, pre-motor/DAN II, cingulo-opercular/action-mode, and auditory, as well as visual networks (V5 and lateral) (Fig. 4c,d; *t*-test, complex *versus* all other networks; *p* < 0.05).

Next, we used overlap scores to generate conjunction maps. For instance, we could generate maps showing all vertices for which SCAN was among the suprathreshold templates, but DMN (dorsolateral component) was not (SCAN+, DMN-) (Fig. 4e, top left). We could also do the opposite, generating maps of vertices labeled as DMN but not SCAN (DMN+, SCAN-) (Fig. 4e, middle left). Of course, then we could also study the intersection, (SCAN+, DMN+; Fig. 4e, bottom left). In the specific case of DMN and SCAN, the amount of co-labeling is near minimal. In contrast, SCAN and pre-motor/DAN II exhibit high levels of overlap–i.e. there are many vertices for which both SCAN and pre-motor/DAN II are among the suprathreshold networks (Fig. 4e, bottom right).

In principle, this procedure can be extended to consider not only pairs of networks, but also triplets, quartets, and so on. Given our focus on SCAN, we identified all vertices for which SCAN was among the suprathreshold networks and the networks with which SCAN was most frequently included as a pair, triplet, and so on (Fig. 4f). We found that the most common pairs were SCAN + SMN foot (33%) and SCAN + SMN face (25%) of roughly 4700 total pairs; the most common triplets were SCAN + SMN foot + SMN hand (28%) and SCAN + pre-motor/DAN II + SMN hand (13%); the most common quartets were SCAN + SMN foot + SMN hand + pre-motor/DAN II (17%) and SCAN + auditory + SMN face + cingulo-opercular/action-mode networks (12%) (see Fig. 4f for quintets and sex-tets). We also calculated a configuration agnostic co-assignment measure–i.e. a simple count of how frequently SCAN appeared with any network, irrespective of whether they formed a pair or were part of a larger complex. We found that SCAN’s most frequent co-labels were pre-motor/DAN II an cingulo-opercular/action-mode (Fig. 4g).

Together these results suggest that SCAN might occupy a hub position by integrating effector, attention, and goal-directed action networks into a shared neural substrate. We note also that SCAN’s overlap with higher-order, association networks was virtually nonexistent, which further reinforces the principle of segregation between sensorimotor and association systems.

### SCAN selectively co-activates with different networks across time

To this point we have considered, exclusively, time-invariant features of SCAN. However, correlations (how we generally measure FC) are a summary of time-varying co-fluctuations. Many studies have shown that (*a*) network activity [18, 25–27, 44–48] (and possibly connectivity) varies over time and (*b*) this variation can be compressed into a small set of “states” – patterns of activity or connectivity that approximately repeat over time. Here, we shift focus from SCAN’s static connectivity onto its time-varying properties. Specifically, we study how activations of SCAN nodes coincide with activations of other networks using “co-activation patterns” or CAPs [44, 49, 50].

To detect SCAN CAPs we developed the following procedure. First, we extracted time series from all SCAN vertices (Fig. 5a). For each imaging session, we calculated the amplitude of SCAN activation as the root mean squared activity these vertices (Fig. 5b) and extracted the amplitude and timing (frame index) of all local maxima. We then repeated this procedure using a temporal null model in which vertex time series were independently circularly shifted by some random offset, exactly preserving their mean/variance and approximately their autocorrelation while destroying inter-vertex correlations. We compared amplitudes of the observed peaks against a null distribution of amplitudes from shuffled peaks and retained only those frames with significantly greater amplitude (p-values for each frame were estimated non-parametrically against 1000 randomizations; we corrected for multiple comparisons by fixing the false discovery rate at *q* = 1% and adjusted the critical *p*-value accordingly; mean*±*standard deviation adjusted critical value across participants of *p*_*adj*_ = 0.0219*±*0.0023). The resulting set of frames represented instants when the collective activity of SCAN is of exceptionally high amplitude. We then extracted whole-brain activity patterns during these frames, concatenated them across imaging sessions, participants, and datasets (Fig. 5c), and clustered them using a k-means algorithm with Lin’s concordance as a measure of similarity (Fig. 5d). The clustering algorithm was repeated 1000 times with random restarts, yielding an ensemble of similar but non-identical partitions. To obtain a consensus partition, we calculated the co-assignment probability of all activation patterns, transformed this to a distance (1 - similarity), and clustered this co-assignment matrix using deterministic agglomerative hierarchical clustering, selecting the optimal number of clusters as the level at which the mean Adjusted Rand Index (ARI; a measure partition similarity) with respect to the ensemble of partitions achieved its maxima.

**Figure 5.**
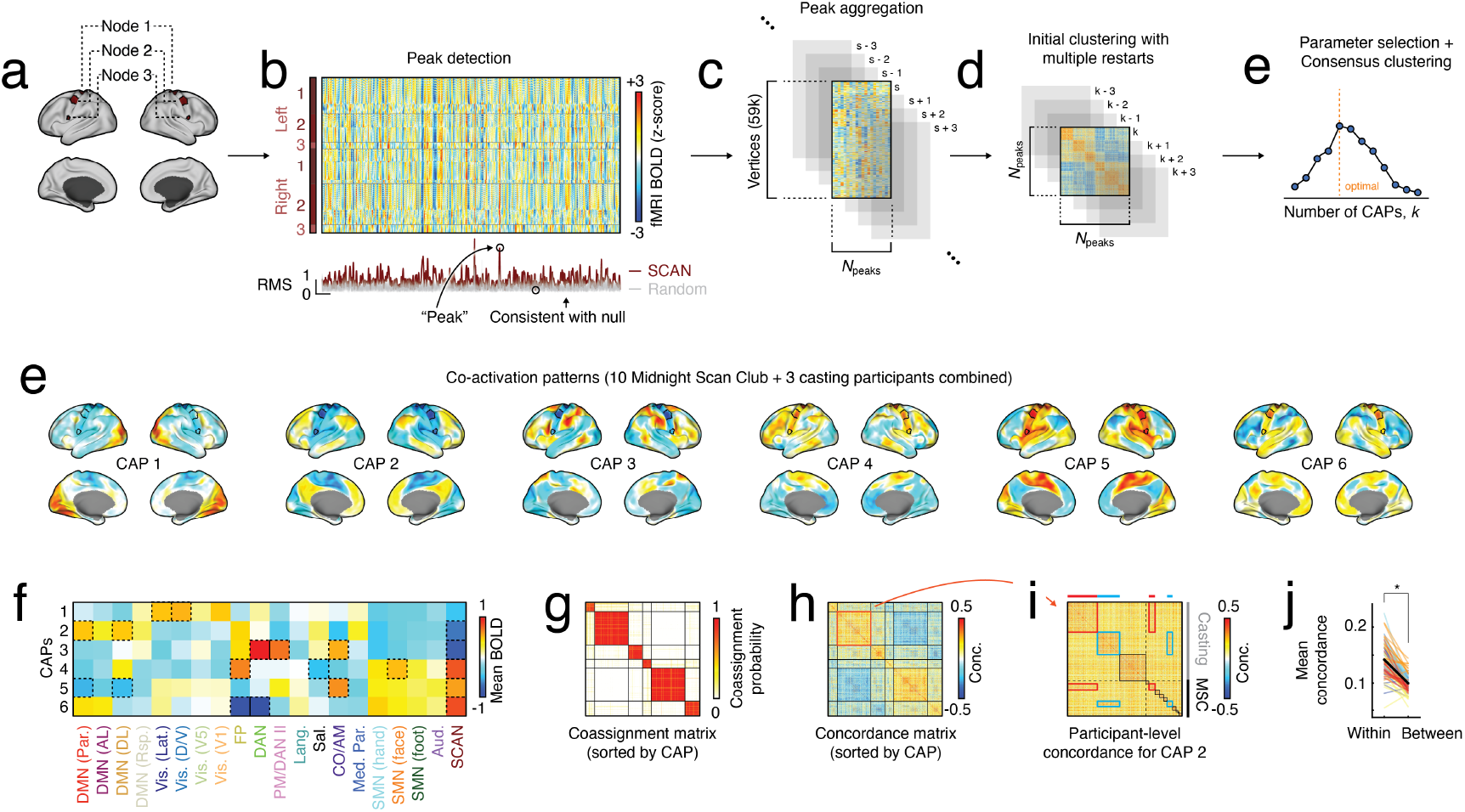
Co-activation pattern (CAP) detection and characterization. Panels *a*-*d* illustrate a schematicized version our pipeline for detecting CAPs. (*a*) First, we extract z-scored fMRI BOLD time series for all SCAN vertices. (*b*) We define SCAN activations as frames when the mean squared activity across all SCAN vertices exceeds that of a temporal null model. We identify all local maxima (peaks) and (*c*) extract the whole-brain activity pattern at those instants. (*c*) We repeat this procedure independently for each imaging session and concatenate all peak activation patterns into a vertex × peak matrix. (*d*) We partition these peaks into clusters based on their similarity to one another. We define CAPs to be the centroids of consensus partitions and interpret the activation patterns assigned to each cluster as a noisy realization of the corresponding CAP. (*e*) Consensus CAPs for pooled MSC data displayed on cortical surfaces. Note that CAPs are organized as anti-correlated pairs (1 + 4, 2 + 5, 3 + 6). (*f*) Similarity matrix (Lin’s concordance) ordered by CAPs. Within each cluster we find evidence that CAPs exhibit additional personalization, (*g*) Inset showing elements that fall within CAP 3 ordered by participant. (*h*) Mean within- and between-participant similarity. Each pair of points represents one of six CAPs. (*i*) Average activity within networks for each of the six CAPs.

We found that at *k* = 6 the clustering algorithm consistently converged to a stable solution. The consensus partition identified six CAPs that could be grouped into three anti-correlated pairs (Fig. 5f; outlined cells correspond to systems whose internal activity exceeded a null distribution under spin tests [51, 52]). For CAPs 1 + 4, SCAN activity was strongest in sub-region 1 and was paired with opposing activation of multiple visual networks (lateral and dorsoventral); for CAPs 2 + 5, SCAN activity was strong across all three sub-regions and was paired with opposed activation of DMN sub-networks (the parietal and dorsolateral components were exceptionally strong); for CAPs 3 + 5, SCAN activity was, again, distributed across all three SCAN sub-nodes and paired with opposed activation of fronto-parietal, dorsal attention, and pre-motor/dorsal attention II networks (Fig. 5f).

As expected, the sorted co-assignment and concordance matrices exhibited strong block diagonal structure (Fig. 5g,h). Interestingly, we observed that clusters exhibited non-trivial sub-structure, wherein within-cluster activation patterns could be further sub-divided based on participant (Fig. 5i,j), suggesting that though CAPs are broadly shared across individuals, they undergo additional refinement at the subject level. Importantly, casting participants 1 and 2 (sub-cast1; sub-cast2) were also participants of in the MSC study (see highlighted red and blue blocks in Fig. 5i). We found that the similarity of CAPs estimated from the casting dataset could be used to identify those individuals in MSC (see red and blue insets in panel Fig. 5i; *t*-test, mean concordance within casting participants sub-cast1 and sub-cast2 versus all other participants; *p* < 0.05). We also calculated a number of potentially useful statistics for SCAN CAPs–e.g. transition probabilities between CAPs as well as differences in their duration, amplitude, and peak velocities (see Fig. S4).

Together, these results suggest that SCAN can be viewed as an impermanent object that co-activates with other networks through a low-dimensional set of patterns. Its strongest co-activations include networks with which it maintains strong static FC–e.g. somatomotor network, pre-motor/DAN II, and cingulo-opercular/action-mode–but also exhibits strong opposed activation with networks to which it maintains weak FC–e.g. fronto-parietal, dorsal attention, and sub-components of the DMN.

### Mapping CAPs to functional connectivity

In the previous sections, we identified a set of CAPs– whole-brain patterns of activity, temporally aligned to high-amplitude activity in SCAN–that recur across time and imaging sessions. CAPs live in activity space. That is, they have dimensions *N*_*vertices*_ × 1 and describe the level of BOLD activity in surface vertices. However, CAPs can be directly mapped to FC, which has dimensions *N*_*vertices*_ × *N*_*vertices*_ by adopting an “edge-centric” approach [29, 30, 53–64]. Specifically, the outer product of any activation pattern, including CAPs, with itself yields a *N*_*vertices*_ × *N*_*vertices*_ matrix that describes the instantaneous magnitude and sign of co-fluctuation between pairs of vertices (see Fig. 6a). Indeed, if we calculated co-fluctuation matrices using BOLD data from each frame of an imaging session, then the temporal average of the corresponding co-fluctuation matrices would equal, exactly, static FC. That is, the co-fluctuation matrices are the time-resolved “building blocks” of FC.

**Figure 6.**
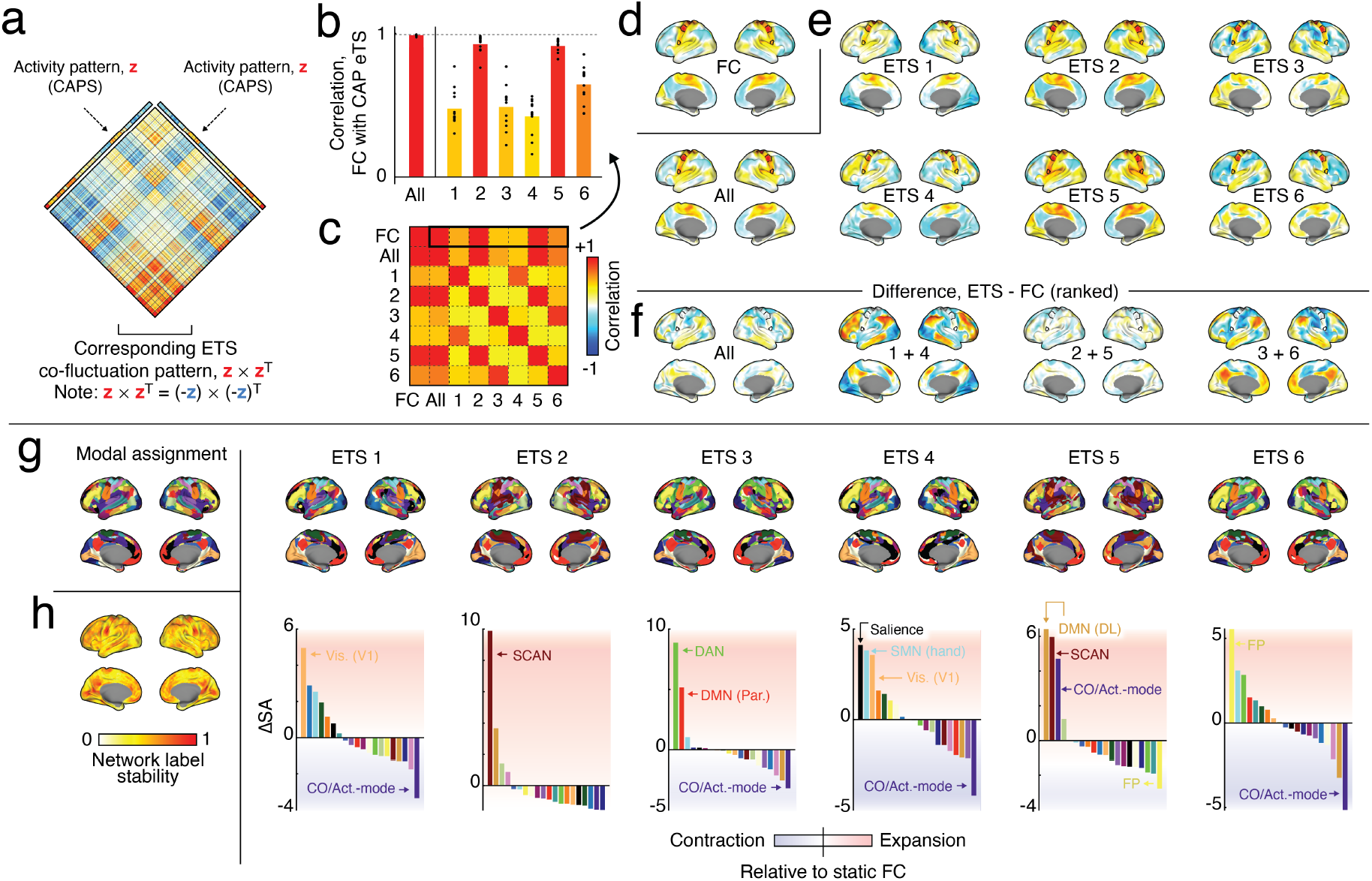
Mapping CAPs to FC. Edge time series serve as a framework for bridging activity and connectivity. Specifically, one can map a vertex × 1 vector to a vertex × vertex co-fluctuation matrix. For a given imaging session, the summation and normalization of co-fluctuation matrices across time is exactly equal to FC. (*a*) A visual explanation depicting how a given CAP can be transformed into a co-fluctuation matrix by taking the outer product with itself. (*b*) Correlation of FC with co-fluctuation matrices associated with each CAP. Each black dot represents an MSC participant. (*d*) SCAN FC seed map. (*e*) SCAN seed maps for each of the six CAPs independently and the seed map estimated after pooling all CAPs into a single meta-cluster. (*f*) Vertex-wise difference between FC and co-fluctuation patterns estimated for each CAP pair. As with static FC, we can use template matching to label vertices for each CAP based on whole-brain co-fluctuation patterns constructed using the procedure illustrated in *a*. When we do this, we can compare the network labels of vertices estimated for each CAP to the network labels estimated at rest, to identify networks with high rates of expansion or contraction in terms of total surface area. (*g*, left) Modal network assignments across all participants estimated using static FC. (*g*, right/top) Network assignments estimated for each of the CAPs following edge time series transformation into co-fluctuation matrices. (*g*, right/bottom) Change in surface area for each network. Large positive valeus correspond to expansion relative to static FC; large negative values correspond to contraction. (*h*) Relative network stability averaged over all CAPs. Values close to 1 correspond to vertices whose network label is consistent across CAPs. Values close to zero are those that change network assignment frequently.

Here, we capitalize on this intuition and construct co-fluctuation matrices for all CAPs, independently, as well as an omnibus co-fluctuation matrix constructed by pooling all peak activations (effectively treating them as a single CAP). With these matrices in hand, we can ask to what extent individual CAPs are aligned with and differentially contribute to SCAN seed FC.

Briefly, we find when CAPs are combined, we obtain a near perfect reconstruction of static FC (mean *r* = 0.98*±*0.03; Fig. 6b,c). This is in line with previous studies showing that high-amplitude frames, at the whole-brain level, explain whole-brain FC above and beyond other categories of frames [58, 65]; the distinction here is that we show an analogous effect at the network level [57]. When we examine individual CAPs, we find that they 1) collapse from six patterns into into three distinct modes of co-fluctuation, 2) each of which maintains a unique relationship with static FC. Specifically, CAPs 1 + 4, 2 + 5, and 3 + 6 exhibit nearly indistinguishable co-fluctuation patterns with respect to one another (Fig. 6c). This pairing effect is a direct consequence of how activations get transformed to connectivity; an activity pattern and itself negated will give rise to the same co-fluctuation matrix. Given that the CAPs were organized into anti-correlated pairs, this effect was anticipated.

This mapping of CAPs to connectivity allows us to not only determine overall similarity to SCAN seed FC, but also to localize where CAP co-fluctuations deviates from SCAN seed FC. To do this, ranked each element of SCAN seed FC and each CAP pair co-fluctuation map. This ranking procedure is necessary to match the range of the data; the FC seed map is bounded between -1 and 1, whereas the CAPs co-fluctuation maps are, in principle, unbounded. Following this transformation, we observe CAP pair-specific effects. As expected, the set of combined CAPs and the 3 + 6 pair (the most frequently observed pair of CAPs and the pair that had the strongest global correspondence with the FC seed map), exhibited only minor local differences with respect to FC. CAP pairs 1 + 4 and 2 + 5, on the other hand, exhibited more extreme differences. In the case of CAP pair 1 + 4, we observed increased connectivity of SCAN to cingulo-opercular/action-mode network and to sub-components of the visual network, alongside reduced connectivity to sub-components of the default mode and frontoparietal networks.

Collectively, these results underscore the relationship between generic high-amplitude activity for explaining static FC–i.e. when pooled together, CAPs co-fluctuations yield a near perfect match to the SCAN FC seed map. However, they also demonstrate that, on a per-CAP basis, there exists some heterogeneity such that their corresponding co-fluctuation patterns do not simply mimic SCAN’s static FC pattern.

### Personalization of CAPs and time-varying connectivity

In the previous sub-sections, we focused largely on group-representative CAPs estimated after pooling MSC and casting data. The cluster centroids – the CAPs – that we obtained from this analysis were therefore composites across individuals. It is unclear whether we would observe similar CAPs if we were to perform the clustering and CAPs detection procedure using only data a single individual–e.g. participants from the casting study.

To address this, we separately clustered SCAN activation patterns for each of the three casting and ten MSC participants. To facilitate comparisons, we forced the number of clusters to *k* = 6 (the optimal number from the group analysis). We then used the Hungarian algorithm to match participant-level CAPs to the CAPs estimated using the group data.

### CAPs rate is linked to motor network plasticity

Over the last several sub-sections, we detected and characterized SCAN CAPs, demonstrating that these patterns may be broadly shared at the population level but subtly refined individually. We converted CAPs from activations to connectivity using edge time series and demonstrated that the detected CAPs correspond to distinct connectivity patterns (though collectively explain static FC almost completely). Further, we showed that CAPs have distinct subcortical and cerebellar components. In this final section, we demonstrate that SCAN CAPs track with experimental stages in the casting study.

As part of the casting study, each participant was imaged prior to, during, and following upper limb immobilization. The aim was to measure plasticity effects of this fairly extreme experimental manipulation (casting one’s own arm for extended periods of time). Here, we seek to link this experimental condition with dynamic properties of SCAN, namely the frequency with which certain CAPs appear. The original casting studies, which focused on FC of somatomotor regions to the rest of the brain and motor disuse pulses, predate the discovery of SCAN. Consequently, and despite the fact that SCAN may play an important role in helping coordinate intentional action, how its properties vary over the course of the casting experiment is unknown.

There are many possible strategies for linking SCAN with the experimental data. We opted for a simple approach. Specifically, we compared CAP rates – the number of times a given CAP appeared within an imaging session, divided by the amount of usable data for that session – across pre-casting, casting, and post-casting conditions. We found that, when all CAPs were combined, there was a tendency (though not significant) for CAPs rate to decrease (`post - pre`; permutation test, *p* = 0.052; Fig. 8). However, when we examined individual CAPs, we found that CAPs 1, 2, and 6 all exhibited statistically significant differences in their rates (maximum *p* = 0.03; Fig. 8). These observations suggest that CAPs frequency – and therefore elements of brain network dynamics – may be sensitive to and modulated by the limb immobilization procedure.

**Figure 7.**
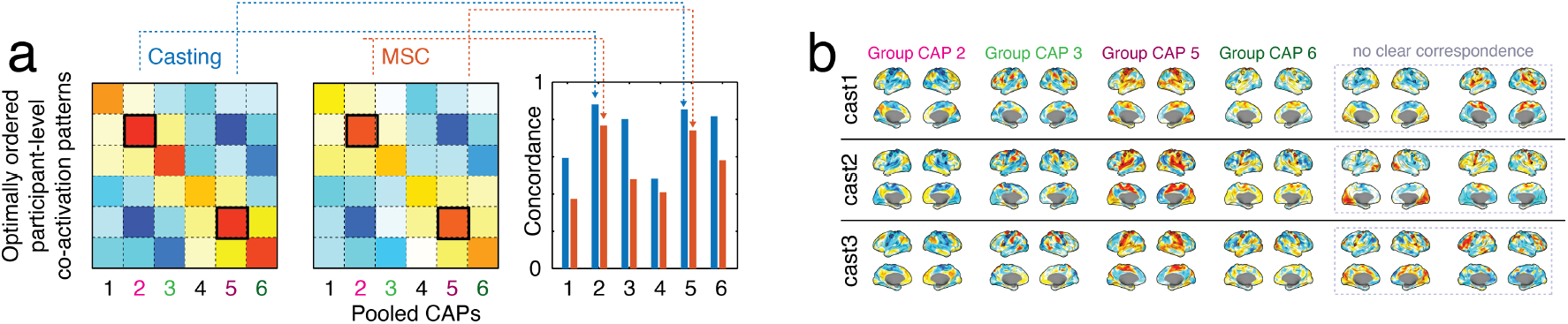
Comparing individualized and group CAPs. We pooled data from all participants to estimate CAPs. Independently, we estimated CAPs for each participant using only their imaging data. We performed this analysis for both the MSC and casting datasets. We used the Hungarian algorithm to obtain the best mapping of individualized to group CAPs. (*a*) Here, we show the similarity matrix with optimized ordering of both casting and MSC CAPs to the group CAPs. The diagonal of these matrices (isolated in the right-most sub-panel) is the focus – it describes how similarity of the matched CAPs to one another. Group CAPs 2 + 5 (and to a lesser extent 3 + 6) appear in single-participant datasets, whereas CAPs 1 + 4 are not especially similar. (*b*) Personalized CAPs from each of the casting participants highlighting the four CAPs with good representation at both the group and individual level, as well as two “outliers” for each participant that had no obvious correspondence.

**Figure 8.**
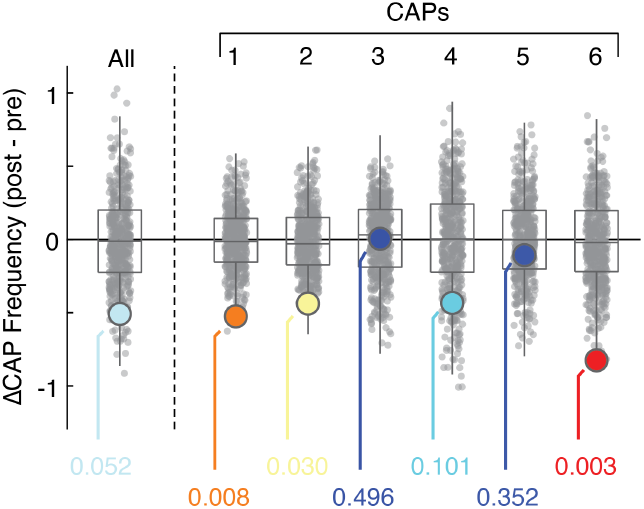
Differences in CAP frequency following limb immobilization. We calculated normalized and standardized CAP frequency for all CAPs together and for each CAP individually. We then calculated the mean difference in CAP frequency for `post-pre` imaging sessions. We then compared this observed difference against a null model in which we, within subjects, circularly shifted the session labels. This circular shift preserves the spatial autocorrelation but otherwise destroys the correspondence with the CAP counts. Negative scores indicate CAPs whose frequency decreased following casting. The colored points are the observed changes in frequency and are colored based on a linear mapping of the corresponding p-value to the colormap (red = small p-values; blue = large). The gray points represent subsets (1000 samples) from the null distribution.

## DISCUSSION

Here, we documented a number of previously undisclosed properties of SCAN. These included static features, such as internal heterogeneity in the connectivity profiles of its constituent vertices and its position in the brain as a hub for network overlap. We also examined SCAN’s time-varying features based on co-activation patterns and transformations of those activations to instantaneous connectivity *via* edge time series. We provided evidence that SCAN activity could be parsimoniously described in terms of six CAPs, broadly organized into three anti-correlated pairs. These CAPs varied in terms of the connectivity profiles, and while one no single CAP fully explained static FC, collectively they did. Finally, we showed that CAP rate – how frequently high-amplitude frames appeared in a given imaging session – decreased following limb immobilization.

### SCAN as a hub of network overlap

One of the most salient features of functional imaging data is that it can be used to partition the cerebral cortex into systems or networks. These networks can be recovered using a variety of techniques, including independent components analysis (ICA) [5], meta-analytic network mapping [66], matching to a set of predefined templates [43, 67], or from data-driven community detection applied to functional connectivity [3, 68–71]. In most cases, the techniques used to estimate community topography explicitly assume a one-to-one mapping of community labels to vertices (though see [55, 72, 73] for examples that explicitly characterize network overlap). That is, each vertex/region/parcel is assigned to one network and one network only.

The assumption of non-overlapping networks is totally reasonable as a first-order approximation of the brain’s functional systems and is line with the perspective that brain function can be localized to specific areas and circuits. However, it necessarily overlooks the possibility that neural elements can belong to multiple systems simultaneously and that they are fundamentally poly-functional or serve as a “gatekeepers” to regulate inter-network information flows [74]. To estimate overlapping networks one generally needs to fit a more complex model. Here, we assess overlap using a variant of template matching, which preserves network labels across individuals (a useful feature for the sake of inter-individual comparison). We find that the entire sensorimotor-attention-action complex has the greatest level of overlap. That is, vertices that make up this complex tend to have strong affiliations with networks past their primary assignment. SCAN, in particular, stands out, as SCAN-primary vertices tend to have affiliations with the greatest number of networks.

On one hand, this observation could reflect the topography of SCAN and its spatial embedding relative to other networks. SCAN is composed of distinct inter-effector regions, each of which maintains borders with multiple networks (see Fig. S3). Network assignments tend to be fuzzy near boundaries, with the probability of a vertex being assigned to a network decaying monotonically and slowly with distance from the border. Combined with the relatively small surface area of the inter-effector regions, one might hypothesize that SCAN’s inter-effector regions are, in essence, positioned at the confluence of many borders. On the other hand, the high levels of overlap could also be a signature of poly-functionality. This is in line with observations from Gordon et al. [11], in which the authors note that SCAN’s connectivity positions it as an intermediary between effectors (somatomotor networks) and networks supporting goal-directed cognition (cinguloopercular/action-mode network).

Future work should investigate the implications and origins of SCAN overlap. For instance, task-fMRI could be used to better understand the conditions under which SCAN is active and the networks with which it tends to co-activate. Is that set large and the experimental conditions diverse? If so, then the high levels of overlap estimated from resting state data takes on additional significance. Indeed, we might hypothesize in that the resting architecture in part constrains the repertoire of task-evoked activity, as previous studies have shown that network boundaries and connectivity features tend to delineate evoked activation patterns [75, 76].

### Stability of SCAN activity and connectivity over time

Functional networks are often described in terms of territories and maps. Maps are relatively easy to visualize and interpret; the set of vertices assigned to a network define an object and its boundaries. The spatial structure of that object – the networks it borders, the connectivity within and between those networks, etc. – can be easily studied from maps of networks. However, this falsely gives the impression that the map is immutable (or at least that the boundaries are meaningful). Many studies have shown that network boundaries can vary over time and across different experimental conditions [77, 78], even within an individual [79]. An even greater number of studies have shown that activity and connectivity change spontaneously and dynamically across time. An important area of research in non-invasive imaging and network neuroscience is to understand the principles by which these brains recon-figure over time.

In this paper, we focused on mapping time-varying changes in SCAN activity and connectivity. We targeted high-amplitude activations, averaged across all inter-effector regions. These frames tend to have stronger signal and are known to contribute disproportionately to static FC [54, 61]. We found that SCAN activations can be grouped into six co-activation patterns or CAPs. We showed that these patterns are mostly preserved at the individual level. Most importantly, these observations suggest that SCAN activations may be low-dimensional and recurrent, in the sense that noisy realizations of the same activation patterns re-occur across time.

Interestingly, when we transformed CAPs into connectivity maps, we found that individually, none of the CAPs fully explained static FC. CAPs 2 and 5 were the most similar in terms of their connectivity. These patterns were also the most frequent, observed in every participant, and recovered at different values of *k* (from *k* = 2 to *k* = 10). However, to fully recapitulate SCAN’s seed connectivity, it was necessary to also include connectivity patterns from the other CAPs (1, 3, 4, and 6), which individually were much less similar to SCAN in terms of connectivity.

Our findings paint a picture of SCAN in which its connectivity is not static, but arises from the superposition of a low-dimensional set of recurring time-varying connectivity patterns. This is viewpoint is not wholly new – e.g. see [58] – and has, at times, been challenged. Specifically, it has been pointed out that many apparently “dynamic” features of BOLD fMRI can be parsimoniously explained by finite sampling from stationary multivariate processes [20, 80, 81]. However, it is unclear whether this “minimally sufficient” model, though capable of reproducing time-varying features of real fMRI BOLD data, has anything to do with the true underlying process by which fMRI BOLD data are generated. The construction and interpretation of model itself involves circularities. Multivariate correlation structure is a summary of the history of linear dependence among a system’s elements; it is not generative. However, when it is treated as generative, it should not be surprising, then, that the outputs it generates – in this case, estimates of time-varying activity and connectivity – resemble the observations used to estimate the correlations.

### Participant-specificity of SCAN dynamics

Historically, estimates of activity and connectivity states have been obtained using large imaging samples like the Human Connectome Project, Adolescent Brain Cognitive Development, and UK Biobank datasets. Due to large numbers of individuals in those datasets, the corresponding state estimates should be viewed as population-level. Increasingly, however, there has been a call for personalized imaging based biomarkers–e.g. precision functional mapping. However, personalized estimates of imaging markers generally require a greater number of observations. The “shallow” datasets described above – lots of brains, but relatively few observations per brain – are not well-suited for this task. Densely-sampled brains in the style of the MSC or the MyConnectome project [82, 83] are better candidates for estimating personalized markers, including brain states.

Here, because we estimated brain states separately at the “group” (though only 11 brains) and single-subject levels, we were able to obtain a crude estimate of how much brain state topography varies from one individual to another and relative to the group. We found that the two most frequent states – CAPs 2 and 5, in which SCAN activation was paired with activation of somatomotor and cingulo-opercular/action-mode networks – were preserved in almost every individual in both the casting and MSC datasets. CAP 5, in partcular, exhibited slightly elevated RMS relative to all other CAPs, suggesting that amplitude (as opposed to pattern) may be a key feature driving the detectability of a given CAP across individuals. CAPs 3 and 6 were evident in casting participants, but less so for MSC. CAPs 1 and 4 were not well recapitulated across participants in either dataset.

These observations suggest that CAPs estimated at the group level should be interpreted cautiously, as pooling and clustering data from across individuals may distort activation profiles, such that the group centroids may reflect blurred composites of many brains but not representative of any individual.

### Future directions

This work presents many avenues for future research. One particularly useful direction is the development of standardized CAPs templates. The same way that template matching is used to generate personalized estimates of network topography, one could create a series of template CAPs from a representative dataset and use those as “lures” – mapping and labeling framewise activation patterns from a secondary, testing dataset to each CAP template. In the same way that template matching atomizes networks, this would atomize CAPs, ensuring rapid and computationally efficient deployment (no need for clustering in a new dataset) and facilitate inter-individual comparisons.

A second important extension of this work involves linking SCAN CAPs with time-varying physiological and behavioral measurements [84]. Implicitly, we expect that CAP transitions reflect meaningful changes in physiology or mental processes or in the parameters of an underlying dynamical system – e.g. changes in excitability or the cumulative effect of training/immobilization. These analyses would simultaneously impact the discussion surrounding the verisimilitude of time-varying fMRI BOLD and connectivity fluctuations. If the fluctuations are robustly linked to meaningful changes in physiology, it suggests that they may be more than simple statistical artifacts.

### Limitations

There are a number of limitations associated with this study. First, we focus on the canonical cortical SCAN network – six inter-effectors organized into three bilaterally symmetric pairs. However, SCAN extends into other cortical areas and includes subcortical and cerebellar components. Here, we ignore these secondary cortical and non-cortical SCAN components, largely as a matter of convenience and in support of a simpler narrative. Future work should examine CAPs and SCAN network overlap beyond the canonical components.

Second, we adopted a specific procedure for detecting CAPs based on selecting high-amplitude frames, estimating an ensemble of partitions using a *k*-means algorithm, and collapsing across ensemble variable using hierarchical clustering to obtain a consensus partition. In principle, there are near infinite variants of this algorithm that could result in diverging outcomes. For instance, rather than clustering high-amplitude frames, we could have tried to cluster *all* frames. Rather than selecting frames with the highest amplitude across all SCAN vertices, we could have isolated frames in which specific SCAN nodes–i.e. specific inter-effectors– exhibited peak activity. The choice of clustering algorithms are also variable; *k*-means is convenient in that it forces a specific number of clusters, but it also forces every frame to belong to its best-fitting cluster, even if that fit is quite poor. Other algorithms might handle these types of outliers more effectively, or allow for clusters to be arranged hierarchically into nested families of CAPs [56, 85]. In summary, future work should explore alternative CAP detection pipelines.

## MATERIALS AND METHODS

### Datasets

#### Midnight Scan Club

The Midnight Scanning Club (MSC) dataset focused on the precise characterization of ten individual subjects via collection of large amounts of per-individual data. Each subject underwent twelve separate two-hour scanning sessions. In the first two sessions, four T1 images, four T2 images, four MR angiograms, and eight MR venograms were collected. In the last ten sessions, five hours of resting-state fMRI data and over five and a half hours of task fMRI data across three different tasks was collected. Participants also underwent extensive neuropsychological testing. Here we analyzed the resting-state data. Here, we analyzed post-processed resting state data (https://openneuro.org/datasets/ds000224/versions/1.0.1).

#### Casting

Three healthy adult participants wore a cast covering the entire right upper extremity for two weeks. They were scanned every day for 6-9 weeks. Scans included 42-64 daily 30-minute resting-state functional MRI scans before, during and after casting. Participants later underwent 12-24 additional scans as part of a control experiment. In all, 27-43 hours of resting-state functional MRI data in each individual were collected. Here, we analyzed post-processed resting state data (https://openneuro.org/datasets/ds002766/versions/3.0.2).

#### Additional post-processing steps

Neither datasets had undergone spatial smoothing as part of the pre-processing procedures. We concatenated dense time series data from all sessions (z-scoring vertex time courses first at the session level and after concatenation at the dataset level) and spatially smoothed vertex time courses (three-dimensional 2.55mm FWHM kernel).

### Network assignments

We used a probabilistic mapping approach to assign network labels to cortical vertices. Specifically, we used vertex-to-network probability maps estimated from [39] (available here: https://github.com/cjl2007/PFM-Depression/tree/main/PFM-Tutorial/Utilities). These maps assign each vertex, *i* ∈ {1, …, 59412}, a probability of belonging to one of 20 canonical resting-state networks, *c* ∈ {1, …, 20}. The full set of maps were arranged in a matrix, **X** *∈* R59412×20. The element *P*_*ic*_ therefore denotes the probability that vertex *i* was assigned to network *c*.

To obtain personalized network labels, we performed the following analysis. Let 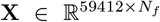 be the vertex-by-frame matrix of BOLD-fMRI time courses, where *N*_*f*_ is the total number of frames and varies with participant and session. Using these maps, we obtained a network-representative time courses as: **X**_*c*_ = **P**^⊺^**X**. The resulting matrix has dimensions 20 × *N*_*f*_ and each row corresponds to a network.

We then computed brain-wide correlation maps by calculating 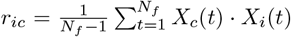 for each vertex, *i*, and network, *c*. We assigned each vertex the network label, *c*, corresponding to the index that maximizes the following expression: max_*c*_ *r*_*ic*_. These labels serve as estimates of personalized network topography.

### Overlapping network labels

The above procedure uses a winner-take-all approach for assigning vertices to networks. Alternatively, we can obtain overlapping estimates of networks by assigning a vertex, *i*, to all networks *c* such that *r*_*ic*_ *> r*_*threshold*_. From this, we can calculate the number of networks to which a vertex is assigned.

### Co-activation patterns

#### Peak detection

One of the aims of the paper was to estimate co-activation patterns (CAPs). That is, detect frames corresponding to high-amplitude SCAN activity and cluster them into distinct groups. In essence, CAPs tries to determine what the rest of the brain is doing in terms of its activity when SCAN is highly active.

To obtain CAPs, we performed the following analyses for each network. First, using participants’ personalized network labels, we estimated the time-varying BOLD signal amplitude for SCAN as: 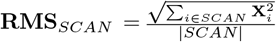. Here, we let **X** = [*X* (1), … *X* (*N*_*f*_)]. Note that, in principle, this approach could be tailored to any network (for example, see [57]).

Next, we obtained all local maxima of **RMS**_*SCAN*_, excluding frames flagged for high levels of in-scanner motion. We then compared the amplitude of these frames against a null model in which we randomly and independently circularly shifted SCAN vertex activation time courses (within imaging sessions), calculated SCAN RMS based on these randomized time courses, and selected the peak RMS values. We repeated this procedure 1000 times, generating a null distribution of peak RMS values. We then compared the observed peaks against this null distribution and calculated a non-parametric *p*-value for each peak as the proportion of the null distribution that exceeded its amplitude. We retained only those frames whose observed RMS was significantly greater than that of the null distribution, correcting for multiple comparisons correction by adjusting in the critical value using the Benjamini-Hochberg procedure (with false discovery rate fixed at *q* = 0.01) [86]).

#### Clustering peak activation maps into CAPs

To cluster peak activation patterns into states, we used a two-stage algorithm. First, we grouped states into clusters using a k-means algorithm with Lin’s concordance as the distance metric [87]. Lin’s concordance is a similarity measure that resolves to the familiar correlation similarity value when the vectors have similar means and variances. Deviations in terms of the mean and variance will necessarily penalize the similarity. Given peak activation patterns, *p* and *q*, we calculate their concordance as:

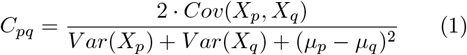

where *Cov*() and *Var*() are the covariance and variance operators, respectively. The variables *X*_*p*_ and *µ*_*p*_ denote the whole-brain activation pattern at peak, *p*, and the mean activity of that pattern, respectively.

We ran the k-means algorithm with 1000 random restarts from *k* = 2 to *k* = 10. In general, each restart yielded a dissimilar cluster solution, allowing us to explore variation in solution. However, we also wanted to obtain a singular point estimate of the cluster solutions. To consolidate across these noisy estimated, we used a consensus clustering approach. Briefly, for a given *k*, we calculated the cluster coassignment matrix, whose elements counted the fraction of 1000 solutions in which a pair of activation patterns were assigned to the same cluster. We transformed these probabilities to a distance (1 - probability), and used a deterministic and agglomerative hierarchical clustering algorithm to partition activation patterns into consensus clusters (with 1 - probability as the distance metric and with the “average” linkage function), selecting the hierarchical level whose consensus solution maximized the adjusted Rand index (ARI) with respect to the ensemble of 1000 partitions generated by k-means [88]. ARI is a partition similarity measure that is bounded to the interval [0, 1], with 0 and 1 corresponding to maximal dissimilarity and similarity, respectively. The resulting

Intuitively, our CAPs clustering approach leverages the stochasticity of the k-means algorithm to generate an initial sample of plausible cluster solutions. We then compress those estimates as a co-assignment matrix and take advantage of the deterministic hierarchical clustering algorithm to obtain a point estimate that represents, in effect, the midpoint of the initial sample–a consensus solution [89].

### Edge time series

Each CAP has dimensions of vertices × 1 and represents a pattern of activity. To transform activations into their corresponding functional connectivity contribution, we used “edge time series.” Edge time series (eTS) are a procedure for temporally unfolding a correlation coefficient into its framewise contributions. The bivariate product-moment correlation is calculated as:

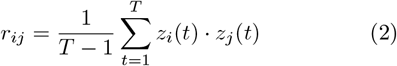

where *T* is the number of frames and *z*_*i*_(*t*) is the z-scored activity of vertex *i* at time *t*. Omitting the summation step yields the following time series:

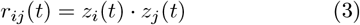

The elements of this time series, if we were to sum them up, return the familiar correlation coefficient, and we can therefore interpret them as the framewise contributions to the correlation. The elements tell us when and in what direction two fMRI BOLD time series are co-fluctuation. That is, *r*_*ij*_(*t*) is positive if the activity of *i* and *j* is concordant (fluctuating in the same direction) and takes on a large value if one (or both) of *i* and *j* is deflecting far from its mean.

We can extend this idea to obtain framewise contributions not just to individual correlation coefficients (functional connections), but to whole-brain connectivity. For instance, if we collected *r*_*ij*_(*t*) for all {*i, j*} pairs, we can arrange those elements in a vertex × vertex matrix. This matrix represents the instantaneous (at time *t*) whole-brain connectivity pattern.

Here, we use edge time series to transform CAPs into their corresponding connectivity contributions. For a given set of activation patterns–e.g. those assigned to CAP *c*–these patterns have dimensions vertex × number of patterns. We simply take product of this array with itself transposed and normalize each entry by the number of patterns to generate a vertex × vertex array. This array represents the connectivity contribution of CAP *c*.

## ACKNOWLEDGMENTS

RFB acknowledges support from the National Science Foundation (NCS-FO award #2023985).

## Data availability statement

Post-processed casting data are available publicly at: https://openneuro.org/datasets/ds002766/versions/3.0.2. Midnight scan club data are also available: https://openneuro.org/datasets/ds000224/versions/1.0.1.

## SUPPLEMENTARY MATERIALS

**Figure S1.**
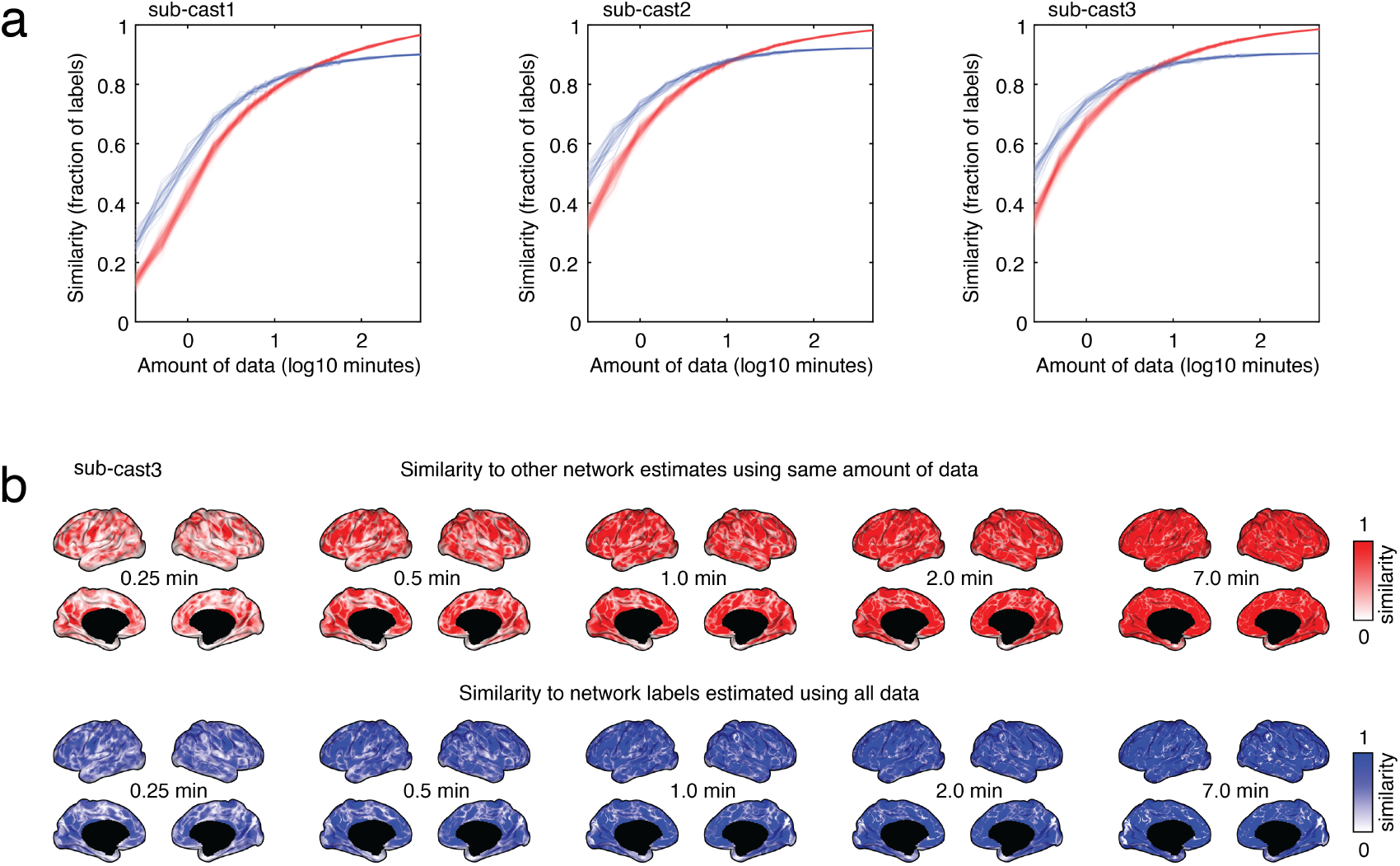
Stability of network labels estimated using temporal template matching for “casting” participants. We subsampled different numbers of frames from each participant’s imaging data. We then used these frames to obtain network estimates. We repeated this procedure multiple times (25 repetitions). We then compared these labels to the network labels estimated using *all* samples and to other network labels estimated using the same number of subsampled frames. (*a*) Similarity (number of vertices with the same label) of sub-sampled network labels relative to each other (red) and to the full sample (blue). (*b*) Same measures but without spatial averaging and depicted in anatomical space.

**Figure S2.**
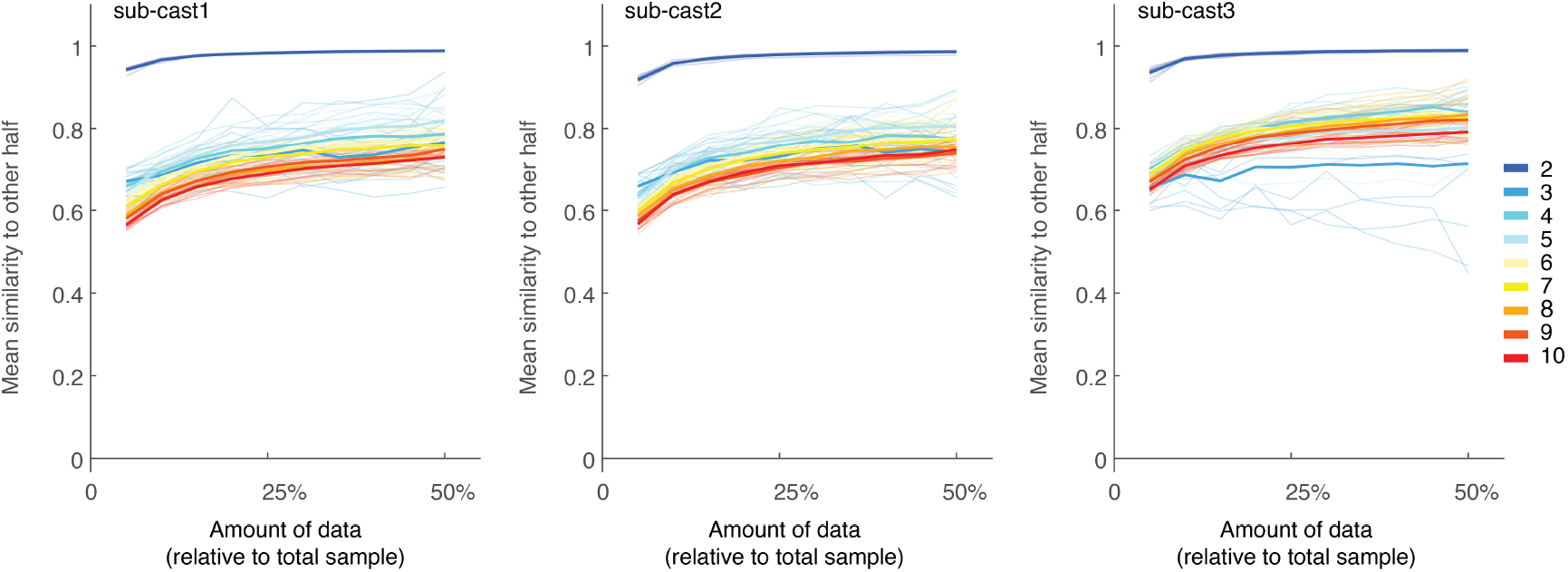
Stability of CAP labels as a function of the amount of data used to estimate CAPs. We split each participants data into two halves. For the first half (training data), we used the full sample to estimate CAPs at *k* = 2, …, 10. For the second half (test data), we subsampled frames in incremements of 10% and estimated CAPs for the subsample. For a given subsample size, we repeated the subsampling and CAPs estimation procedure 10 times. We then used the Hungarian algorithm to match CAPs to the training data and calculated the mean spatial similarity across all of the maps. Values near 1 indicate that the CAPs detected in the test data could be matched to those estimated in the training data. Values closer to 0 indicate poor alignment of test-training CAPs. We found that with *k* = 2, CAPs could almost always be reliably estimated on a per-subject basis, even with small amounts of data. However, there was a big gap in the ability to recover CAPs for *k >* 2. This gap narrowed as more data was included, but never saturated. These observations suggest that accurate characterization of CAPs at participant level may require prohibitively large amounts of data. We note, however, that these results are contingent on the specific algorithms used to detect high-amplitude frames and to cluster them into putative CAPs.

**Figure S3.**
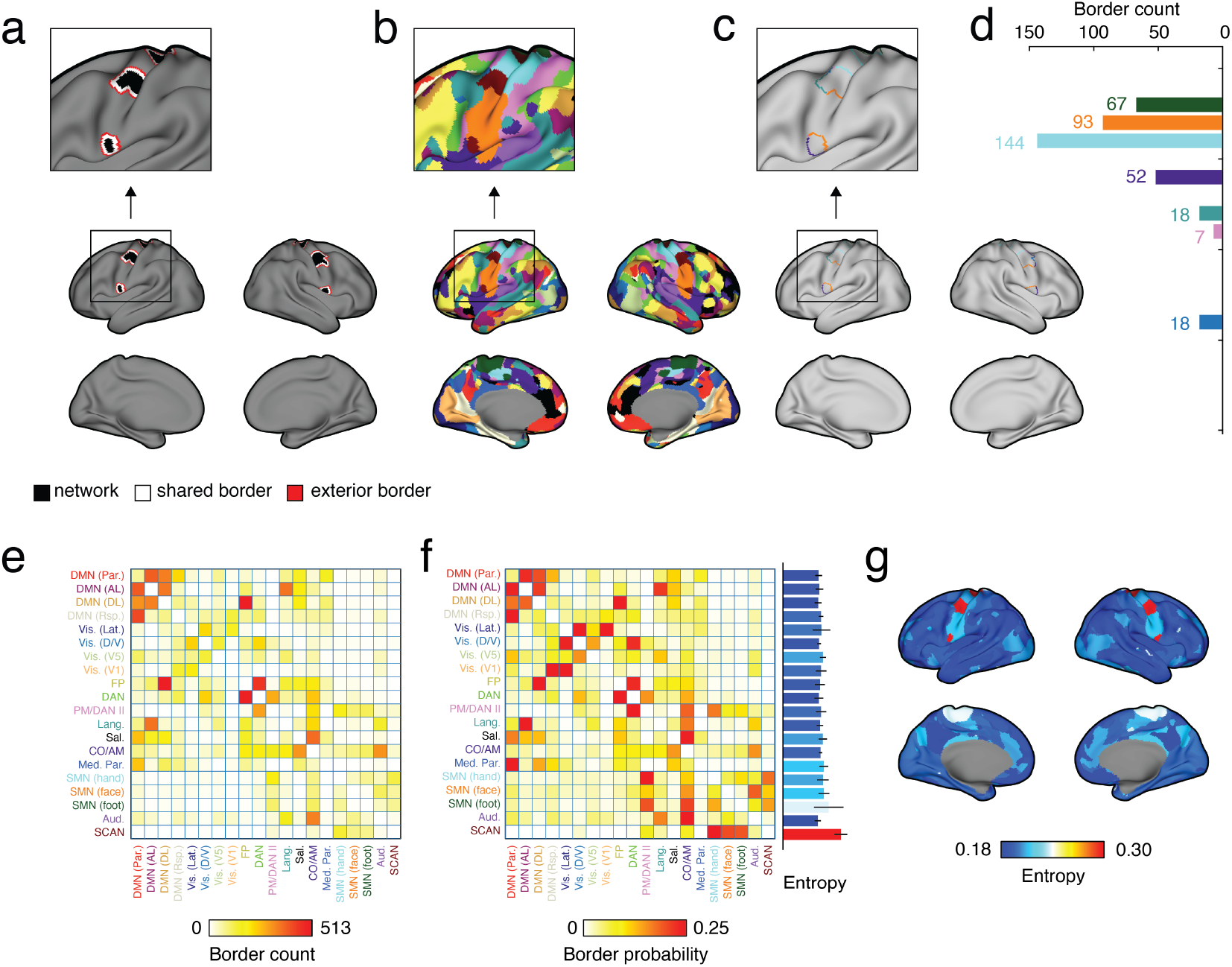
Border statistics. We examined the border statistics around SCAN inter-effector regions and all other networks to assess whether these statistics might explain why SCAN exhibited high levels of network overlap. Specifically, we identified (*a*) the border around each networks as the vertices. Briefly and for a given network, we identified all faces that included at least one vertex assigned to that network. We defined the border as vertices that formed a face with this set of vertices. We referred to these vertices as forming the “shared border.” We then examined all vertices that formed a face with at least one of the shared border vertices. We referred to this set of vertices as the “exterior border” (see panel *a*). In parallel, we had network labels for each vertex (see *b*). (*c*) We used the exterior border to mask the network labels and calculated the frequency with which other networks appeared in this set (see *d*). (*e*) We repeated this procedure for all networks, generating a matrix of how frequently each of the 20 networks appear along the border of all other networks. (*f*) We can transform these counts to probabilities by normalizing each row to have a sum of 1. We can also calculate the entropy of each row – i.e. for a given network how much uncertainty/variability there was in its border networks (see *f*, right; error bars are estimated across participants). (*g*) Entropy scores plotted on the brain surface.

**Figure S4.**
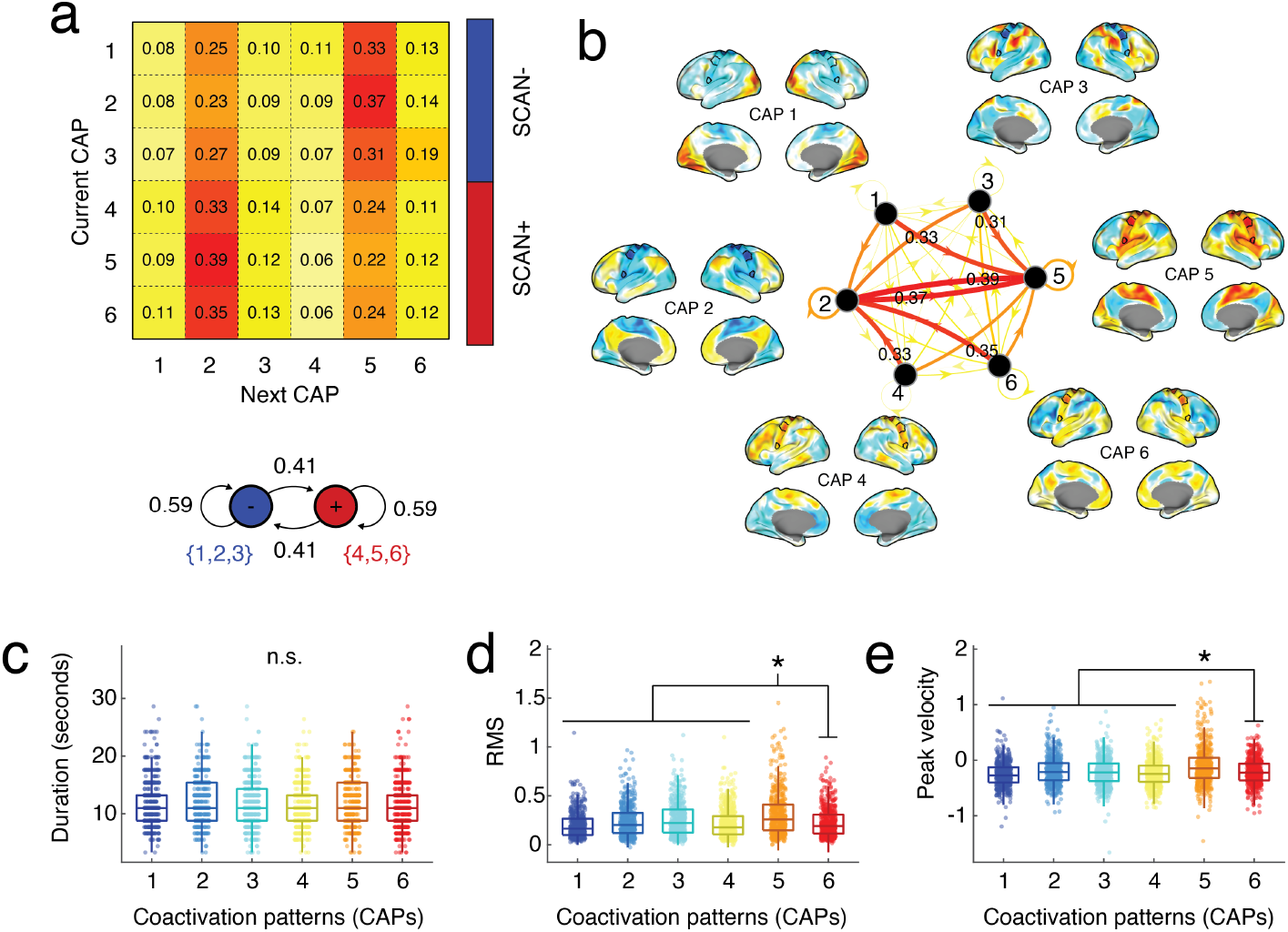
CAP statistics. (*a*) CAP transition matrix. We calculated how frequently one CAP transitioned to another. Because we defined CAPs based on peak activations (which are necessarily separated by *at least* 1 frame, but generally much longer), our transition matrix does not exhibit strong auto-transitions. That is, it is unlikely that two successive peaks are assigned to the same CAP. Our analysis also suggests that SCAN CAPs tend to alternate based on the sign of SCAN activity. That is, if SCAN activity is positive/negative, then the following CAP will tend to be negative/positive (opposite sign). (*b*) We depict the transition matrix graphically and label some of the strongest transitions. We also calculated several statistics for each CAP. Specifically, we calculated (*c*) trough-to-trough duration (number of seconds from the trough immediately preceding a peak to the trough immediately following it); (*d*) CAP RMS; (*e*) and peak velocity – the greatest absolute rate of change in RMS during a CAP’s corresponding trough-to-trough segment. We found no differences in duration across CAPs. However, we found that RMS and peak velocity of CAP 5 – activation of SCAN + cingulo-opercular/action-mode network – was significantly greater than the other CAPs (*t*-tests; both *p* < 10^*−*^5).

## References

[1] Jonathan D Power, Alexander L Cohen, Steven M Nelson, Gagan S Wig, Kelly Anne Barnes, Jessica A Church, Alecia C Vogel, Timothy O Laumann, Fran M Miezin, Bradley L Schlaggar, et al. Functional network organization of the human brain. Neuron, 72(4):665– 678, 2011.

[2] BT Thomas Yeo, Fenna M Krienen, Jorge Sepulcre, Mert R Sabuncu, Danial Lashkari, Marisa Hollinshead, Joshua L Roffman, Jordan W Smoller, Lilla Zöllei, Jonathan R Polimeni, et al. The organization of the human cerebral cortex estimated by intrinsic functional connectivity. Journal of neurophysiology, 2011.

[3] Olaf Sporns and Richard F Betzel. Modular brain networks. Annual review of psychology, 67:613–640, 2016.

[4] Richard F Betzel, John D Medaglia, Ari E Kahn, Jonathan Soffer, Daniel R Schonhaut, and Danielle S Bassett. Structural, geometric and genetic factors predict interregional brain connectivity patterns probed by electrocorticography. Nature biomedical engineering, 3 (11):902–916, 2019.

[5] Stephen M Smith, Peter T Fox, Karla L Miller, David C Glahn, P Mickle Fox, Clare E Mackay, Nicola Filippini, Kate E Watkins, Roberto Toro, Angela R Laird, et al. Correspondence of the brain’s functional architecture during activation and rest. Proceedings of the national academy of sciences, 106(31):13040–13045, 2009.

[6] Constantin Freiherr von Economo and Georg N Koskinas. Die cytoarchitektonik der hirnrinde des erwachsenen menschen. J. Springer, 1925.

[7] Justin L Vincent, Gaurav H Patel, Michael D Fox, Abraham Z Snyder, Justin T Baker, David C Van Essen, John M Zempel, Lawrence H Snyder, Maurizio Corbetta, and Marcus E Raichle. Intrinsic functional architecture in the anaesthetized monkey brain. Nature, 447(7140):83–86, 2007.

[8] Janine Bijsterbosch, Samuel J Harrison, Saad Jbabdi, Mark Woolrich, Christian Beckmann, Stephen Smith, and Eugene P Duff. Challenges and future directions for representations of functional brain organization. Nature neuroscience, 23(12):1484–1495, 2020.

[9] Lucina Q Uddin, Richard F Betzel, Jessica R Cohen, Jessica S Damoiseaux, Felipe De Brigard, Simon B Eickhoff, Alex Fornito, Caterina Gratton, Evan M Gordon, Angela R Laird, et al. Controversies and progress on standardization of large-scale brain network nomenclature. Network Neuroscience, 7(3):864–905, 2023.

[10] Ru Kong, R Nathan Spreng, Aihuiping Xue, Richard F Betzel, Jessica R Cohen, Jessica S Damoiseaux, Felipe De Brigard, Simon B Eickhoff, Alex Fornito, Caterina Gratton, et al. A network correspondence toolbox for quantitative evaluation of novel neuroimaging results. Nature communications, 16(1):2930, 2025.

[11] Evan M Gordon, Roselyne J Chauvin, Andrew N Van, Aishwarya Rajesh, Ashley Nielsen, Dillan J Newbold, Charles J Lynch, Nicole A Seider, Samuel R Krimmel, Kristen M Scheidter, et al. A somato-cognitive action network alternates with effector regions in motor cortex. Nature, 617(7960):351–359, 2023.

[12] Wilder Penfield and Edwin Boldrey. Somatic motor and sensory representation in the cerebral cortex of man as studied by electrical stimulation. Brain: A journal of neurology, 1937.

[13] Nico UF Dosenbach, Marcus Raichle, and Evan M Gordon. The brain’s cingulo-opercular action-mode network. Preprint at PsyArXiv 10.31234/osf.io/2vt79, 2024.

[14] Georgios P Skandalakis, Luca Viganò, Clemens Neudorfer, Marco Rossi, Luca Fornia, Gabriella Cerri, Kelsey P Kinsman, Zabiullah Bajouri, Armin D Tavakkoli, Christos Koutsarnakis, et al. White matter connections within the central sulcus subserving the somatocognitive action network. Brain, page awaf022, 2025.

[15] Jiao Li, Dajing Wang, Jie Xia, Chao Zhang, Yao Meng, Shuo Xu, Huafu Chen, and Wei Liao. Divergent suicidal symptomatic activations converge on somato-cognitive action network in depression. Molecular Psychiatry, 29 (7):1980–1989, 2024.

[16] Jianxun Ren, Wei Zhang, Louisa Dahmani, Qingyu Hu, Changqing Jiang, Yan Bai, Gong-Jun Ji, Ying Zhou, Ping Zhang, Weiwei Wang, et al. The somato-cognitive action network links diverse neuromodulatory targets for parkinson’s disease. bioRxiv, pages 2023–12, 2023.

[17] Gong-Jun Ji, Michael D Fox, Mae Morton-Dutton, Yingru Wang, Jinmei Sun, Panpan Hu, Xingui Chen, Yubao Jiang, Chunyan Zhu, Yanghua Tian, et al. A generalized epilepsy network derived from brain abnormalities and deep brain stimulation. Nature communications, 16(1):2783, 2025.

[18] Daniel J Lurie, Daniel Kessler, Danielle S Bassett, Richard F Betzel, Michael Breakspear, Shella Kheilholz, Aaron Kucyi, Raphaël Liégeois, Martin A Lindquist, Anthony Randal McIntosh, et al. Questions and controversies in the study of time-varying functional connectivity in resting fmri. Network Neuroscience, 4(1):30–69, 2020.

[19] Timothy O Laumann and Abraham Z Snyder. Brain activity is not only for thinking. Current Opinion in Behavioral Sciences, 40:130–136, 2021.

[20] Timothy O Laumann, Abraham Z Snyder, and Caterina Gratton. Challenges in the measurement and interpretation of dynamic functional connectivity. Imaging Neuroscience, 2:1–19, 2024.

[21] Richard F Betzel, Lisa Byrge, Farnaz Zamani Esfahlani, and Daniel P Kennedy. Temporal fluctuations in the brain’s modular architecture during movie-watching. NeuroImage, page 116687, 2020.

[22] R Matthew Hutchison, Thilo Womelsdorf, Elena A Allen, Peter A Bandettini, Vince D Calhoun, Maurizio Corbetta, Stefania Della Penna, Jeff H Duyn, Gary H Glover, Javier Gonzalez-Castillo, et al. Dynamic functional connectivity: promise, issues, and interpretations. Neuroimage, 80:360–378, 2013.

[23] Rikkert Hindriks, Mohit H Adhikari, Yusuke Murayama, Marco Ganzetti, Dante Mantini, Nikos K Logothetis, and Gustavo Deco. Can sliding-window correlations reveal dynamic functional connectivity in restingstate fmri? Neuroimage, 127:242–256, 2016.

[24] Aaron Kucyi and Karen D Davis. Dynamic functional connectivity of the default mode network tracks day-dreaming. Neuroimage, 100:471–480, 2014.

[25] Danielle S Bassett, Nicholas F Wymbs, Mason A Porter, Peter J Mucha, Jean M Carlson, and Scott T Grafton. Dynamic reconfiguration of human brain networks during learning. Proceedings of the National Academy of Sciences, 108(18):7641–7646, 2011.

[26] Taylor Bolt, Jason S Nomi, Danilo Bzdok, Catie Chang, BT Thomas Yeo, Lucina Q Uddin, and Shella D Keilholz. Large-scale intrinsic functional brain organization emerges from three canonical spatiotemporal patterns. bioRxiv, 2021.

[27] Richard F Betzel, Makoto Fukushima, Ye He, Xi-Nian Zuo, and Olaf Sporns. Dynamic fluctuations coincide with periods of high and low modularity in resting-state functional brain networks. NeuroImage, 127:287–297, 2016.

[28] Sarah A Cutts, Evgeny J Chumin, Richard F Betzel, and Olaf Sporns. Temporal variability of brain– behavior relationships in fine-scale dynamics of edge time series. Imaging Neuroscience, 3:imag_a_00443, 2025.

[29] Farnaz Zamani Esfahlani, Lisa Byrge, Jacob Tanner, Olaf Sporns, Daniel Kennedy, and Richard Betzel. Edge-centric analysis of time-varying functional brain networks with applications in autism spectrum disorder. bioRxiv, 2021.

[30] Sarah Greenwell, Joshua Faskowitz, Laura Pritschet, Tyler Santander, Emily G Jacobs, and Richard F Betzel. High-amplitude network co-fluctuations linked to variation in hormone concentrations over menstrual cycle. bioRxiv, 2021.

[31] Richard F Betzel, Theodore D Satterthwaite, Joshua I Gold, and Danielle S Bassett. Positive affect, surprise, and fatigue are correlates of network flexibility. Scientific Reports, 7(1):1–10, 2017.

[32] Gorka Zamora-López, Changsong Zhou, and Jürgen Kurths. Cortical hubs form a module for multisensory integration on top of the hierarchy of cortical networks. Frontiers in neuroinformatics, 4:1, 2010.

[33] Martijn P Van Den Heuvel and Olaf Sporns. Rich-club organization of the human connectome. Journal of Neuroscience, 31(44):15775–15786, 2011.

[34] Evan M Gordon, Timothy O Laumann, Adrian W Gilmore, Dillan J Newbold, Deanna J Greene, Jeffrey J Berg, Mario Ortega, Catherine Hoyt-Drazen, Caterina Gratton, Haoxin Sun, et al. Precision functional mapping of individual human brains. Neuron, 95(4):791– 807, 2017.

[35] Dillan J Newbold, Evan M Gordon, Timothy O Laumann, David F Montez, Nicole A Seider, Sarah J Gross, Annie Zheng, Ashley N Nielsen, Catherine R Hoyt, Jackie M Hampton, et al. Cingulo-opercular control network supports disused motor circuits in standby mode. bioRxiv, 2020.

[36] Dillan J Newbold, Timothy O Laumann, Catherine R Hoyt, Jacqueline M Hampton, David F Montez, Ryan V Raut, Mario Ortega, Anish Mitra, Ashley N Nielsen, Derek B Miller, et al. Plasticity and spontaneous activity pulses in disused human brain circuits. Neuron, 107 (3):580–589, 2020.

[37] Roselyne J Chauvin, Dillan J Newbold, Ashley N Nielsen, Ryland L Miller, Samuel R Krimmel, Athanasia Metoki, Anxu Wang, Andrew N Van, David F Montez, Scott Marek, et al. Disuse-driven plasticity in the human thalamus and putamen. Cell reports, 44(4), 2025.

[38] David C Van Essen, Stephen M Smith, Deanna M Barch, Timothy EJ Behrens, Essa Yacoub, Kamil Ugurbil, Wu-Minn HCP Consortium, et al. The wu-minn human connectome project: an overview. Neuroimage, 80:62–79, 2013.

[39] Charles J Lynch, Immanuel G Elbau, Tommy Ng, Aliza Ayaz, Shasha Zhu, Danielle Wolk, Nicola Manfredi, Megan Johnson, Megan Chang, Jolin Chou, et al. Frontostriatal salience network expansion in individuals in depression. Nature, 633(8030):624–633, 2024.

[40] Evan M Gordon, Timothy O Laumann, Babatunde Adeyemo, Jeremy F Huckins, William M Kelley, and Steven E Petersen. Generation and evaluation of a cortical area parcellation from resting-state correlations. Cerebral cortex, 26(1):288–303, 2016.

[41] Bharat B Biswal and Lucina Q Uddin. The history and future of resting-state functional magnetic resonance imaging. Nature, 641(8065):1121–1131, 2025.

[42] Robert JM Hermosillo, Lucille A Moore, Eric Feczko, Óscar Miranda-Domínguez, Adam Pines, Ally Dworetsky, Gregory Conan, Michael A Mooney, Anita Randolph, Alice Graham, et al. A precision functional atlas of personalized network topography and probabilities. Nature neuroscience, 27(5):1000–1013, 2024.

[43] Evan M Gordon, Timothy O Laumann, Babatunde Adeyemo, and Steven E Petersen. Individual variability of the system-level organization of the human brain. Cerebral cortex, 27(1):386–399, 2017.

[44] Xiao Liu and Jeff H Duyn. Time-varying functional network information extracted from brief instances of spontaneous brain activity. Proceedings of the National Academy of Sciences, 110(11):4392–4397, 2013.

[45] James M Shine, Michael Breakspear, Peter T Bell, Kaylena A Ehgoetz Martens, Richard Shine, Oluwasanmi Koyejo, Olaf Sporns, and Russell A Poldrack. Human cognition involves the dynamic integration of neural activity and neuromodulatory systems. Nature neuroscience, 22(2):289–296, 2019.

[46] Xiaolong Peng, Qi Liu, Catherine S Hubbard, Danhong Wang, Wenzhen Zhu, Michael D Fox, and Hesheng Liu. Robust dynamic brain coactivation states estimated in individuals. Science Advances, 9(3):eabq8566, 2023.

[47] Zirui Huang, Jun Zhang, Jinsong Wu, George A Mashour, and Anthony G Hudetz. Temporal circuit of macroscale dynamic brain activity supports human consciousness. Science advances, 6(11):eaaz0087, 2020.

[48] Erik SB van Oort, Maarten Mennes, Tobias Navarro Schröder, Vinod J Kumar, Nestor I Zaragoza Jimenez, Wolfgang Grodd, Christian F Doeller, and Christian F Beckmann. Human brain parcellation using time courses of instantaneous connectivity. arXiv preprint 1609.04636, 2016.

[49] Fikret Işik Karahanoğlu and Dimitri Van De Ville. Transient brain activity disentangles fmri resting-state dynamics in terms of spatially and temporally overlapping networks. Nature communications, 6(1):1–10, 2015.

[50] Xiao Liu, Catie Chang, and Jeff H Duyn. Decomposition of spontaneous brain activity into distinct fmri coactivation patterns. Frontiers in systems neuroscience, 7:101, 2013.

[51] František Váša and Bratislav Mišić. Null models in network neuroscience. Nature Reviews Neuroscience, 23(8):493–504, 2022.

[52] Ross D Markello and Bratislav Misic. Comparing spatial null models for brain maps. NeuroImage, page 118052, 2021.

[53] Richard F Betzel, Joshua Faskowitz, and Olaf Sporns. Living on the edge: network neuroscience beyond nodes. Trends in cognitive sciences, 27(11):1068–1084, 2023.

[54] Farnaz Zamani Esfahlani, Youngheun Jo, Joshua Faskowitz, Lisa Byrge, Daniel Kennedy, Olaf Sporns, and Richard Betzel. High-amplitude co-fluctuations in cortical activity drive functional connectivity. Proceedings of the National Academy of Sciences, 2020.

[55] Joshua Faskowitz, Farnaz Zamani Esfahlani, Youngheun Jo, Olaf Sporns, and Richard F Betzel. Edge-centric functional network representations of human cerebral cortex reveal overlapping system-level architecture. Nature neuroscience, 23(12):1644–1654, 2020.

[56] Richard Betzel, Sarah Cutts, Jaco Tanner, Sarah Greenwell, Thomas Varley, Joshua Faskowitz, and Olaf Sporns. Hierarchical organization of spontaneous cofluctuations in densely-sampled individuals using fmri. bioRxiv, 2022.

[57] Richard F Betzel, Evgeny Chumin, Farnaz Zamani Esfahlani, Jacob Tanner, and Joshua Faskowitz. System-level high-amplitude co-fluctuations. BioRxiv, pages 2022–07, 2022.

[58] Richard F Betzel, Sarah A Cutts, Sarah Greenwell, Joshua Faskowitz, and Olaf Sporns. Individualized event structure drives individual differences in whole-brain functional connectivity. NeuroImage, 252:118993, 2022.

[59] Maria Pope, Makoto Fukushima, Richard Betzel, and Olaf Sporns. Modular origins of high-amplitude co-fluctuations in fine-scale functional connectivity dynamics. bioRxiv, 2021.

[60] Sarah A Cutts, Joshua Faskowitz, Richard F Betzel, and Olaf Sporns. Uncovering individual differences in fine-scale dynamics of functional connectivity. Cerebral Cortex, 2022.

[61] Olaf Sporns, Joshua Faskowitz, Sofia Teixera, and Richard Betzel. Dynamic expression of brain functional systems disclosed by fine-scale analysis of edge time series. bioRxiv, 2020.

[62] Evgeny J Chumin, Joshua Faskowitz, Farnaz Zamani Esfahlani, Youngheun Jo, Haily Merritt, Jacob Tanner, Sarah A Cutts, Maria Pope, Richard Betzel, and Olaf Sporns. Cortico-subcortical interactions in overlapping communities of edge functional connectivity. NeuroImage, 250:118971, 2022.

[63] Jacob C Tanner, Joshua Faskowitz, Lisa Byrge, Daniel Kennedy, Olaf Sporns, and Richard Betzel. Synchronous high-amplitude co-fluctuations of functional brain networks during movie-watching. bioRxiv, pages 2022–06, 2022.

[64] Haily Merritt, Amanda Mejia, and Richard Betzel. The dual interpretation of edge time series: Time-varying connectivity versus statistical interaction. bioRxiv, pages 2024–08, 2024.

[65] Elisabeth Ragone, Jacob Tanner, Youngheun Jo, Farnaz Zamani Esfahlani, Joshua Faskowitz, Maria Pope, Ludovico Coletta, Alessandro Gozzi, and Richard Betzel. Modular subgraphs in large-scale connectomes underpin spontaneous co-fluctuation” events” in mouse and human brains. bioRxiv, pages 2023–05, 2023.

[66] Nicolas A Crossley, Andrea Mechelli, Petra E Vértes, Toby T Winton-Brown, Ameera X Patel, Cedric E Ginestet, Philip McGuire, and Edward T Bullmore. Cognitive relevance of the community structure of the human brain functional coactivation network. Proceedings of the National Academy of Sciences, 110(28): 11583–11588, 2013.

[67] Caterina Gratton, Timothy O Laumann, Ashley N Nielsen, Deanna J Greene, Evan M Gordon, Adrian W Gilmore, Steven M Nelson, Rebecca S Coalson, Abraham Z Snyder, Bradley L Schlaggar, et al. Functional brain networks are dominated by stable group and individual factors, not cognitive or daily variation. Neuron, 98(2):439–452, 2018.

[68] David Meunier, Renaud Lambiotte, Alex Fornito, Karen Ersche, and Edward T Bullmore. Hierarchical modularity in human brain functional networks. Frontiers in neuroinformatics, 3:37, 2009.

[69] David Meunier, Renaud Lambiotte, and Edward T Bullmore. Modular and hierarchically modular organization of brain networks. Frontiers in neuroscience, 4:200, 2010.

[70] Richard F Betzel. Community detection in network neuroscience. arXiv preprint 2011.06723, 2020.

[71] Farnaz Zamani Esfahlani, Youngheun Jo, Maria Grazia Puxeddu, Haily Merritt, Jacob C Tanner, Sarah Greenwell, Riya Patel, Joshua Faskowitz, and Richard F Betzel. Modularity maximization as a flexible and generic framework for brain network exploratory analysis. arXiv preprint 2106.15428, 2021.

[72] Luiz Pessoa. Beyond disjoint brain networks: overlapping networks for cognition and emotion. Behavioral and Brain Sciences, 39, 2016.

[73] BT Thomas Yeo, Fenna M Krienen, Michael WL Chee, and Randy L Buckner. Estimates of segregation and overlap of functional connectivity networks in the human cerebral cortex. Neuroimage, 88:212–227, 2014.

[74] Caio Seguin, Maria Grazia Puxeddu, Joshua Faskowitz, Richard F Betzel, and Olaf Sporns. Connectome architecture favours within-module diffusion and between-module routing. bioRxiv, pages 2025–02, 2025.

[75] M Fiona Molloy, Zeynep M Saygin, and David E Osher. Predicting high-level visual areas in the absence of task fmri. Scientific reports, 14(1):11376, 2024.

[76] Alexander D Cohen, Ziyi Chen, Oiwi Parker Jones, Chen Niu, and Yang Wang. Regression-based machinelearning approaches to predict task activation using resting-state fmri. Human brain mapping, 41(3):815– 826, 2020.

[77] Lei Wu, Arvind Caprihan, and Vince Calhoun. Tracking spatial dynamics of functional connectivity during a task. NeuroImage, 239:118310, 2021.

[78] Armin Iraji, Zening Fu, Eswar Damaraju, Thomas P DeRamus, Noah Lewis, Juan R Bustillo, Rhoshel K Lenroot, Aysneil Belger, Judith M Ford, Sarah McEwen, et al. Spatial dynamics within and between brain functional domains: A hierarchical approach to study time-varying brain function. Human brain mapping, 40(6):1969–1986, 2019.

[79] Mehraveh Salehi, Abigail S Greene, Amin Karbasi, Xilin Shen, Dustin Scheinost, and R Todd Constable. There is no single functional atlas even for a single individual: Functional parcel definitions change with task. NeuroImage, 208:116366, 2020.

[80] Timothy O Laumann, Abraham Z Snyder, Anish Mitra, Evan M Gordon, Caterina Gratton, Babatunde Adeyemo, Adrian W Gilmore, Steven M Nelson, Jeff J Berg, Deanna J Greene, et al. On the stability of bold fmri correlations. Cerebral cortex, 27(10):4719–4732, 2017.

[81] Zach Ladwig, Benjamin A Seitzman, Ally Dworetsky, Yuhua Yu, Babatunde Adeyemo, Derek M Smith, Steven E Petersen, and Caterina Gratton. Bold cofluctuation ‘events’ are predicted from static functional connectivity. NeuroImage, 260:119476, 2022.

[82] Russell A Poldrack. Precision neuroscience: Dense sampling of individual brains. Neuron, 95(4):727–729, 2017.

[83] Timothy O Laumann, Evan M Gordon, Babatunde Adeyemo, Abraham Z Snyder, Sung Jun Joo, Mei-Yen Chen, Adrian W Gilmore, Kathleen B McDermott, Steven M Nelson, Nico UF Dosenbach, et al. Functional system and areal organization of a highly sampled individual human brain. Neuron, 87(3):657–670, 2015.

[84] Taylor Bolt, Shiyu Wang, Jason S Nomi, Roni Setton, Benjamin P Gold, Blaise deB. Frederick, BT Thomas Yeo, J Jean Chen, Dante Picchioni, Jeff H Duyn, et al. Autonomic physiological coupling of the global fmri signal. Nature Neuroscience, pages 1–9, 2025.

[85] Haily Merritt, Mary Kate Koch, Youngheun Jo, Evgeny J Chumin, and Richard F Betzel. Social ‘envirotyping’the abcd study contextualizes dissociable brain organization and diverging outcomes. bioRxiv, pages 2024–08, 2024.

[86] Yoav Benjamini and Yosef Hochberg. Controlling the false discovery rate: a practical and powerful approach to multiple testing. Journal of the Royal statistical society: series B (Methodological), 57(1):289–300, 1995.

[87] I Lawrence and Kuei Lin. A concordance correlation coefficient to evaluate reproducibility. Biometrics, pages 255–268, 1989.

[88] Amanda L Traud, Eric D Kelsic, Peter J Mucha, and Mason A Porter. Comparing community structure to characteristics in online collegiate social networks. SIAM review, 53(3):526–543, 2011.

[89] Alexander Strehl and Joydeep Ghosh. Cluster ensembles—a knowledge reuse framework for combining multiple partitions. Journal of machine learning research, 3(Dec):583–617, 2002.

